# On the generality and cognitive basis of base-rate neglect

**DOI:** 10.1101/2021.03.11.434913

**Authors:** Elina Stengård, Peter Juslin, Ulrike Hahn, Ronald van den Berg

## Abstract

Base rate neglect refers to people’s apparent tendency to underweight or even ignore base rate information when estimating posterior probabilities for events, such as the probability that a person with a positive cancer-test outcome actually does have cancer. While often replicated, almost all evidence for the phenomenon comes from studies that used problems with extremely low base rates, high hit rates, and low false alarm rates. It is currently unclear whether the effect generalizes to reasoning problems outside this “corner” of the entire problem space. Another limitation of previous studies is that they have focused on describing empirical patterns of the effect at the group level and not so much on the underlying strategies and individual differences. Here, we address these two limitations by testing participants on a broader problem space and modelling their responses at a single-participant level. We find that the empirical patterns that have served as evidence for base-rate neglect generalize to a larger problem space, albeit with large individual differences in the extent with which participants “neglect” base rates. In particular, we find a bi-modal distribution consisting of one group of participants who almost entirely ignore the base rate and another group who almost entirely account for it. This heterogeneity is reflected in the cognitive modeling results: participants in the former group were best captured by a linear-additive model, while participants in the latter group were best captured by a Bayesian model. We find little evidence for heuristic models. Altogether, these results suggest that the effect known as “base-rate neglect” generalizes to a large set of reasoning problems, but varies largely across participants and may need a reinterpretation in terms of the underlying cognitive mechanisms.

## 1. Introduction

The question of whether the human mind adheres to the rules of probability theory has been debated ever since probability theory itself was developed a couple of hundred years ago. Since then the view has shifted drastically from Laplace’s idea that probability theory is, in essence, common sense reduced to calculus (Brookes et al., 1953) to Kahneman’s and Tversky’s claim that people are unable to follow the rules of probability and instead have to rely on simple heuristics which often lead to fairly accurate answers but at other times produce large biases (Kahneman & Tversky, 1973). A more recent suggestion is that people use an adaptive toolbox of fast and frugal heuristics (Gigerenzer & Todd, 1999) that takes advantage of information in real environments to make accurate (ecologically rational) inferences, applicable also in states of uncertainty and incomplete knowledge, where “optimization” is often not computable.

A phenomenon that has inspired, and been used to exemplify, this research on heuristics and biases is *base-rate neglect*, people’s tendency to respond to the evidence immediately at hand, while ignoring the base-rate (or: prior) probability of an event. Although the original context of base-rate neglect emphasized that it is caused by people’s reliance on simplifying heuristics (Kahneman & Tversky, 1973), the explanation and interpretation of the phenomenon remains debated to this day. Another approach emphasizes that people address base-rate problems much like any other multiple-cue judgment task (Brehmer, 1994; Karelaia et al., 2008). On this view, people typically have a qualitative understanding that both the evidence and base-rate is relevant in base-rate problems, but they typically add up these cues, rather than multiply them as prescribed by probability theory (Juslin et al., 2009). A third proposal is that the underlying information integration is in fact consistent with probability theory, but corrupted by random noise in the process (e.g., Costello & Watts, 2014) or dampened with prior beliefs (Sanborn & Chater, 2016), which appears as base-rate neglect in the empirical data.

### 1.1 Base-rate neglect

One task that often has been used to argue both for and against rationality in human probabilistic reasoning strategies, is the medical diagnosis task. Since the introduction of this task (Casscells et al., 1978) it has been formulated in many different ways. A typical example is the following:

> Suppose that 0.1% of all people in a population carry a virus. A diagnostic test for this virus detects it in 100% of the people who have the virus, but also gives “false alarms” on 5% of the people who do not have the virus. What is the chance that a person with a positive test result actually has the virus, assuming you know nothing about the person’s symptoms or signs?

The correct answer can be computed using Bayes’ theorem and gives a probability of ∼2%. In the formulation above, the problem is specified in a “normalized” format, meaning that the information about the base rate (prevalence), hit rate (probability that a carrier of the virus is tested positive), and false alarm rate (probability that a non-carrier of the virus is tested positive) is given as percentages or single-event probabilities. People tend to overestimate the correct answer substantially, presumably due to putting too much weight on the hit rate and false alarm rate, while largely ignoring the base rate – a phenomenon commonly referred to as *base-rate neglect* or the *base rate fallacy* (Meehl & Rosen, 1955) (see Koehler 1996 for a review and a critical discussion of the phenomenon).

Evidence for base-rate neglect has been found in numerous studies, using different variations of Bayesian inference tasks (see Bar-Hillel, 1980; Barbey & Sloman, 2007; Kahneman & Tversky, 1973, for a few highly influential examples). The level of neglect varies, but the number of correct responses is seldom above 20% (see McDowell & Jacobs, 2017 for a meta-analysis of 35 studies). A complicating factor in explaining the effect is that its magnitude depends on the structure of the task. A number of facilitating factors has been explored that increase participants’ use of base rates in Bayesian inference tasks, such as manipulating the base rate within subjects (Ajzen, 1977; Birnbaum & Mellers, 1983; Fischhoff et al., 1979), emphasizing the relevance of the base rate by highlighting a causal link to the inference task (Ajzen, 1977; Bar-Hillel, 1980; Fishbein, 2015), by providing explicit feedback, and by training (Goodie & Fantino, 1999). What these manipulations have in common is that they make decision makers more sensitive to base rates.

Although base-rate neglect is a well-established fallacy in the decision making literature, there are also studies that have found that people do respond to both the base rate and the hit rate, and instead are neglecting false alarm rates (Juslin et al., 2011). This relates to the tendency of people to be influenced by diagnostically irrelevant information and disregarding relevant information (but see also Crupi et al., 2009), a phenomenon known as “pseudo-diagnosticity” (Ofir, 1988). In the medical diagnosis task, this could be manifested as being influenced by a high hit rate without taking a high false-alarm rate sufficiently into account.

There are also findings suggesting that the severity of base-rate neglect depends on the numerical format of the presented information. In particular, people are better able to reason in accordance with Bayes’ rule when all information is given in terms of *naturally sampled frequencies* (e.g. “95 out of 100 tested people” instead of “95%”)^1^, possibly because cognitive algorithms have evolved to compute with counts and not with probabilities or percentages (Gigerenzer & Hoffrage, 1995). The medical diagnosis problem above can be translated to a natural frequency format as follows:

> Suppose that one person in a population of 1,000 people carries a particular virus. A diagnostic test for this virus gives a positive test result on the person carrying the virus as well as for 50 of the 999 healthy persons. What is the chance that a person with a positive test result actually has the virus, assuming you know nothing about the person’s symptoms or signs?

Gigerenzer and Hoffrage found that in this format the proportion of correct answers increased to approximately 50% compared to 16% with the normalized format. The reason why a natural frequency format is beneficial is still debated. Besides the proposal by Gigerenzer and Hoffrage, there is a dual-process theory (Evans & Stanovich, 2013; Sloman, 1996) that proposes that natural frequencies make people shift from using a primitive associative judgment system to a deliberate rule-based system (Barbey & Sloman, 2007; Sloman et al., 2003).

In addition to using different numerical formats in Bayesian inference tasks, some studies have also used different visual formats to present the relevant information, for example by using Venn diagrams to represent normalized frequencies or collections of dots to represent counts. Adding a pictorial representation of the information has in some cases been shown to enhance participant’s performance (Brase, 2009; Garcia-Retamero & Hoffrage, 2013). However, the results are mixed and it is clear that not all visual representations are helpful (Khan et al., 2015). Importantly, pictorial representations can also be used as a way of providing probability information to participants without giving them exact numbers (see, e.g., Harris, Corner, & Hahn, 2009; Harris, De Molière, Soh, & Hahn, 2017). The sense of uncertainty that this produces may make the experimental paradigm more representative for human reasoning in natural environments, where knowledge about base rates, hit rates, and false alarm rates is rarely exact (Juslin et al., 2009).

### 1.2 Explanations of base-rate neglect

Although the phenomenon of base-rate neglect has been known since at least the 1950s (Meehl & Rosen, 1955) the psychological explanation behind it is still a subject of discussion.

#### 1.2.1 Representativeness heuristic

A first theory was put forth by Kahneman and Tversky, who suggested that the phenomenon is caused by people relying on the representativeness heuristic (Kahneman & Tversky, 1973). They used a task in which participants were presented with personality descriptions of people drawn from a population with a known proportion of lawyers and engineers. Based on the personality descriptions the participants had to predict if the randomly drawn individual was a lawyer or an engineer. People often seemed to disregard the base rate proportions of lawyers and engineers and to base their predictions only on the personality descriptions. Kahneman and Tversky’s explanation was that people assess the representativeness (or similarity) of the personal description to the prototypical member of the professional categories (e.g., of a lawyer). Similarly, in the medical diagnosis task the probability assessment could be based on how representative a positive test outcome is for a diseased versus a healthy person. If it is considered more representative for a diseased person then the probability that the person has the virus is predicted to be high. Although there have been attempts to formulate the representativeness heuristic into a computational model (Bordalo et al., 2020; Dougherty et al., 1999; Juslin & Persson, 2002; Nilsson et al., 2008), its application to the sort of base-rate task considered here has not been examined. A problem with the representative heuristic is that it predicts that base rates are always ignored entirely. However, many empirical findings suggest that it is not an all-or-nothing phenomenon, but can differ in severity based on moderating factors, such as the format in which the problem is presented. While the representativeness heuristic can possibly account for base-rate neglect in some tasks, it is unclear how it would account for moderating factors.

#### 1.2.2 Heuristic toolbox

Gigerenzer and colleagues claim that as long as information is presented in the natural frequency format, people often make the appropriate computations and do not commit the base rate fallacy (Gigerenzer & Hoffrage, 1995). If, however, the information is presented in a normalized format, people will rely on heuristics, such as reporting the hit rate or the difference between the hit rate and the false alarm rate, some of which will lead to base-rate neglect effects. The effects of moderating factors on base-rate neglect can, in principle, be accounted for by shifts between different heuristics, although the specific mechanism for choice between heuristics in the base-rate task remain unspecified (but see Rieskamp & Otto, 2006 and Marewski & Schooler, 2011 for models of selection of heuristics in related domains).

#### 1.2.3 Linear-additive integration

In sequential belief revision tasks it has long been known that rather than relying on Bayesian integration people tend to average the “old” and the “new” data (Hogarth & Einhorn, 1992; Lopes, 1985; Shanteau, 1975). This work suggests that people have a tendency to combine information in a linear-additive fashion even when multiplicative integration is the normative solution, which is a model that has also found support in more recent work (Juslin et al., 2008, 2011). On this view, biases are due to people using strategies that are well adapted to cognitive constraints and constraints of a noisy real-life environment. In the context of the medical diagnosis task, this means that the participants may understand that the base rate and the hit rate should be used, but without sufficient appreciation for the functional form implied by Bayes’ theorem, they integrate them in an additive rather than multiplicative manner. Base-rate neglect arises when a participant assigns too little weight to the base rate in comparison to the optimal weight (i.e., the weight that produces the best linear-additive approximation of the Bayesian responses). The moderating factors on base-rate neglect are accounted for by people using various contextual cues to determine the weighting of the base-, hit-, and false alarm rate. The linear additive account of base-rate neglect is both indirectly supported by the literature on multiple-cue judgment (Brehmer, 1994) and directly by the results from computational modeling on base-rate problems (Juslin et al., 2011). It resembles the Bayesian model in the sense that it assumes that participants appreciate that multiple cues should be considered and that they should be integrated in a manner that puts higher weight on more informative cues. It is, however, ‘heuristic’ in its integration stage, where the (non-linear) Bayesian rule is replaced by a simple (linear) rule.

#### 1.2.4 Explanations based on probability theory

Another class of explanations of base-rate neglect is based on probability theory. This class consists of two conceptually very different models. The first is based on the idea that cognitive judgments are corrupted by random error (Erev et al., 1994; Hilbert, 2012). Based on this idea, Costello and Watts (Costello & Watts, 2014, 2016, 2017, 2018, 2019) proposed the “Probability theory plus noise” model, which produces regression-like effects that may make it look as if base rates are being neglected. The effects of moderating factors can to some extent be accounted for by changes in the magnitude of the random noise. The other type is based on the idea that people use Bayesian judgement strategies that incorporate prior beliefs. In this model, regressive effects are largely the result of the “dampening” of the responses, caused by the prior. A recent example is the “Bayesian sampler” model (e.g., Sanborn & Chater, 2016; Zhu, Sanborn, & Chater, 2020). The effects of moderating factors can be accounted for by differences in the strength of prior beliefs.

### 1.3 Limitations in current literature

While previous studies have provided key insights into how humans reason with probabilities, they have also left many questions unanswered.

Firstly, many of the previous studies have only examined problems similar to the ones used in the examples above, that is, problems with an extremely low base rate, a hit rate close or equal to one, and a false alarm rate close to zero (e.g., Khan et al., 2015; Sloman et al., 2003). Although there are studies that have tested participants on more than one trial (e.g., Fischhoff et al., 1979; Gigerenzer & Hoffrage, 1995; Juslin et al., 2011), none so far have performed a systematic exploration of the space of possible stimulus values. Consequently, it remains unknown how representative the results obtained in a rather extreme “corner” of this space are for human reasoning in general.

Secondly, since many studies used only one trial per participant, hardly any modeling has been performed at the level of individual participants. An exception is Juslin et al. (2011), where it was reported that most participants used a linear additive integration to approximate Bayes theorem, both when naively addressing the base-rate problems and after explicit instruction on Bayes theorem. Little is however currently known about individual judgment strategies.

Lastly, biological information processing is constrained by various factors, such as neural noise (Faisal et al., 2008), the costliness of neural computation (Lennie, 2003), and limits on the precision with which neural systems can approximate optimal solutions (Beck et al., 2012). The imperfections caused by these limitations can be modelled as “decision noise”, which is believed to be a major source of errors in perception (e.g., Drugowitsch, Wyart, Devauchelle,

& Koechlin, 2016; Stengard & Van Den Berg, 2019) and may potentially also explain biases in cognition (Erev et al., 1994; Hilbert, 2012). At present, models based on probability theory are the only ones incorporating a form of noise. To properly compare competing theories about cognitive strategies, they need to be equalized in terms of their noise assumptions.

### 1.4 Purpose of the present study

To address these limitations, we tested participants on a broad range of cases of the medical diagnosis task, in which we systematically varied the base rate, hit rate, and false alarm rate. The participants were tested on one out of four tasks, which differed in the frequency format of the provided information (normalized vs. natural frequencies) and the visual presentation format (symbolic vs. pictorial). Finally, we used a cognitive modelling approach to try gain insight into the cognitive strategies employed by the participants and the role of noise in accounting for their responses.

## 2. Methods

### 2.1 Data sharing

The data are available at https://osf.io/3vkad/. Modeling code will be made available upon publication of the paper.

### 2.2 Participants

Forty lab participants (31 females, 9 males; mean age 25.6 years, age span 20-45 years) were recruited from the student population at the Department of Psychology at Uppsala University and 289 online participants were recruited on the crowd-sourcing service Amazon Mechanical Turk (MTurk). The lab participants were compensated with cinema tickets or gift vouchers with a value equivalent to approximately $10 per hour. The qualification requirement for participating in the study on MTurk were a Human Intelligence Task (HIT) approval rate greater than 98% and Number of HITs approved greater than 5000. The online participants in the control experiment were compensated $0.30 for approximately 2 minutes of work and online participants in the main experiment were compensated $5 for approximately 30 minutes of work.

### 2.3 Materials and Design

#### 2.3.1 Control experiment

To verify that our paradigm replicates the original findings of base-rate neglect, we tested 100 of the online participants on a task with a single question, formulated in the same way as in many earlier experiments: “A new disease that is transmitted via a virus has been discovered. Researchers have developed a test that indicates whether a person has the virus or not. A person who has the virus will always get a positive test result. However, the test is not perfect and sometimes a perfectly healthy person still gets a positive test result. 0.1% of people have the virus. Among those who don’t have the virus there is a 5% chance of getting an incorrect positive test result. Imagine meeting a randomly chosen person who has gotten a positive test result. What is the probability that this person actually has the virus, assuming that you know nothing about their health status?”. Participants gave their answer as a percentage, typed in a text field. They could give the answer in decimal precision but all of them provided an integer answer.

#### 2.3.2 Main experiment

On every trial, participants received information about the base rate of a fictitious virus in a fictitious hospital. They were also informed about the hit- and false-alarm rates of a medical test designed to detect the existence of the virus. The task was to estimate the probability that a randomly chosen person from the hospital who had received a positive test result actually had the virus. Participants provided their answer in percentages in all of the conditions. In the symbolic tasks the response was given by typing the numbers on a keyboard and in the pictorial tasks by clicking on a number line. Within each participant, we factorially crossed five base rates (0.1, 0.3, 0.5, 0.7, 0.9), three hit rates (0.5, 0.7, 0.9), and three false alarm rates (0.1, 0.3, 0.5), resulting in a total of 45 trials. Note that the stimulus values were restricted to the “sensible” part of the stimulus space in which the hit rates were at least 50% and false alarm rates were at most 50%. Between participants, we factorially crossed two frequency formats (“natural frequency” and “normalized frequency”) and two visual presentation formats (“symbolic” and “pictorial”).

The two conditions with symbolic presentation format (Figure 1A-B) were performed by 189 online participants. These conditions were similar to how information was presented to participants in most previous studies using the medical diagnosis task. In this presentation format, base rates, hit rates, and false alarm rates were all presented numerically. Seven participants were excluded because they did not complete the whole experiment. All analyses were performed on the data from the remaining 182 participants (65 female, 115 male, 2 other; mean age 34.1 years, age span 19-70).

**Figure 1.**
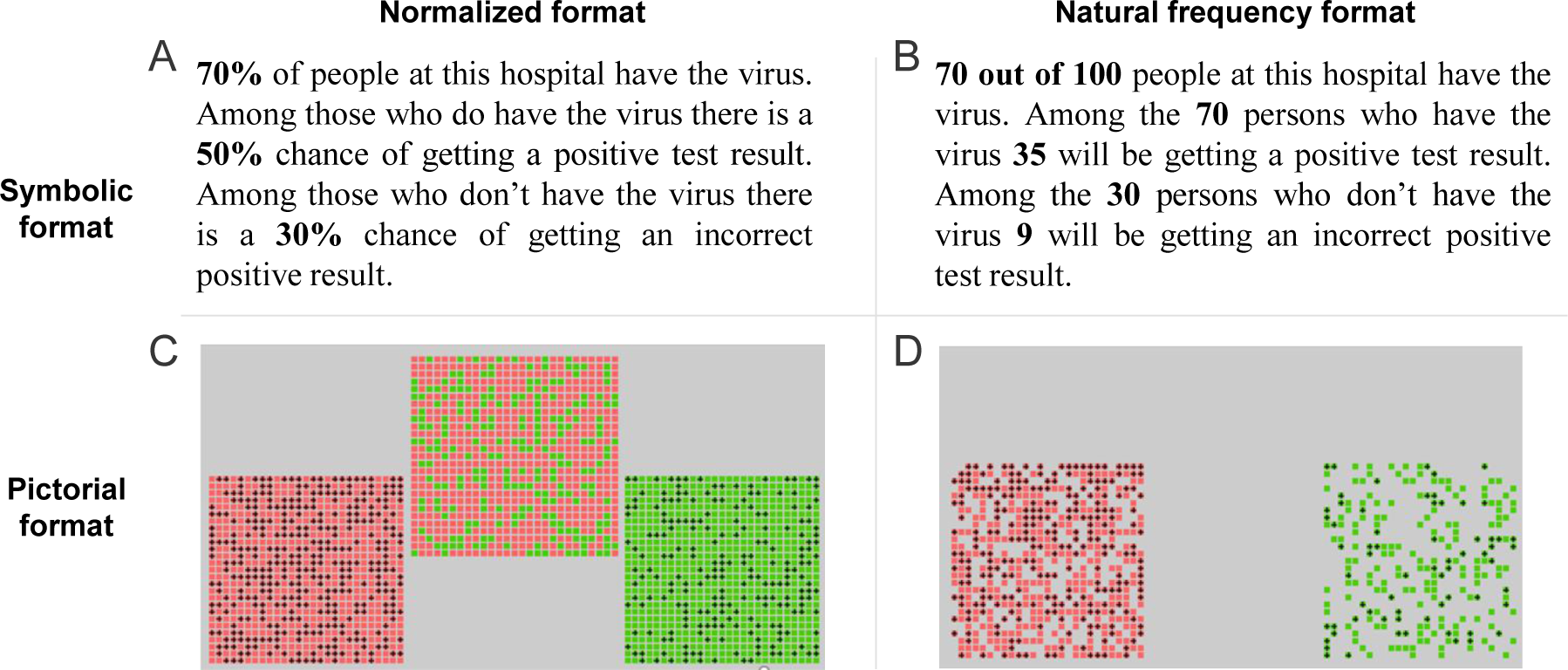
Experimental design. Illustration of one trial in each of the four conditions. Top left: symbolic normalized format. Top right: symbolic natural frequency format. Bottom left: pictorial normalized format. Bottom right: pictorial natural frequency format. The pictorial examples are screenshots from the actual experiment, while the symbolic examples are translations of the original stimuli (which were presented in Swedish).

The two conditions with the pictorial presentation format (Figure 1C-D) were performed by the 40 lab participants. In these conditions, the information was represented by means of “probability matrices” similar to the ones used by Harris et al. (Harris et al., 2009, 2017). In these matrices, every single square represents a person. The colour of the square signaled whether the person had the virus (red) or not (green) and the presence of a plus sign represented a positive result on the medical test. Each matrix consisted of 27 by 27 squares, which presumably was large enough to discourage participants from explicitly counting them. Stimuli were generated using the Psychophysics Toolbox (Brainard, 1997) for Matlab. In the tasks with normalized format, the participants were shown three matrices, separately representing the base rate, hit rate, and false alarm rate (Figure 1C). For the natural-frequency format, all the information could in principle be presented within a single matrix, but to increase visibility the hit rate and the false alarm rates were separated on the screen (Figure 1D).

### 2.4 Procedure

#### 2.4.1 Symbolic task conditions

Data for these two conditions were collected online, using the Amazon Turk platform. The task consisted of 45 trials in which information was presented either as natural frequencies (91 participants; Figure 1A) or as proportions (91 participants; Figure 1B). We randomized the trial sequence and then kept it the same for all participants. The starting point in this sequence varied between participants, in such a way that every test item was presented as the first trial for at least two participants. The participants in these conditions received general information about the experiment and gave informed consent before starting the task. They were encouraged to make their best judgments and performed the 45 trials in a self-paced manner. They were also informed that they were not allowed to use any kind of calculator. Completion time varied strongly across participants, from 5 to 76 minutes (*Mdn* = 16 min).

#### 2.4.2 Pictorial task conditions

The conditions with pictorial stimulus presentations were conducted in the lab. At the start of the session, the participant received general information about the experiment and gave informed consent. Thereafter, the experimenter left the room and the participant would start the experiment. Each participant performed the experiment with information presented to them either in natural frequency format (20 participants; Figure 1C) or in normalized format (20 participants; Figure 1D). The same 45 items as in the symbolic task were used, but now presented twice and with the trial order randomized per participant. Participants performed the experiment in a self-paced manner. Completion times varied from 6 to 69 minutes (*Mdn* = 17 min).

#### 2.4.3 Discrimination task

After the main task, the participants in the pictorial conditions performed an additional discrimination task with 200 trials in which they were shown a single matrix (identical to the ones used in the main task) and were asked to estimate the proportion of squares that had plus signs on them. This task was used to assess the level of noise in the participant’s estimations of the base rates, hit rates, and false alarm rates and was later used to put a constraint on the model parameters in the main task.

### 2.5 Computational modeling

We fit four models that represent the three generic theories in regard to the cognitive processes that underlie these judgments. Heuristic Toolbox Theory claims that people forego part of the information and avoid integration of the normative cues (Gigerenzer & Hoffrage, 1995). Linear additive models claim that, even if people have learned to appreciate the relevance of base rates and hit rates, they lack insight about the functional form of Bayes’ theorem, and – as in other multiple-cue judgment tasks – they default to additive cue integration (Juslin et al., 2009). The third theory is that the cognitive processes integrate the cues according to Bayes’ theorem, but there is random noise in the process that appears as biases in data (Costello & Watts, 2014). The first theory emphasizes heuristic *selection* of cues, the second theory emphasizes heuristic *integration* of cues, and the third theory highlights normative but noisy integration of the cues. While it would be naïve to expect the fitting of these models to 48 responses to precisely identify the exact parameters used by each individual participant, as illustrated in Figures A1 and A5 in the Appendix, the models make clearly distinguishable predictions. The results can therefore be informative with regard to the kind cognitive of strategies used. For example, evidence in favor of the Bayesian and linear additive models over the heuristic models suggests that people do integrate the cues into a posterior probability. Likewise, evidence for the Bayesian over the linear additive model that the integration respects the functional form of Bayes’ theorem.

Each model implements a decision process that takes the base rate (BR), hit rate (HR) and false-alarm rate (FAR) as input and maps this to a predicted response, *R*. This mapping consists of two stages (Figure 2). First, a deterministic integration rule is applied to map the input triplet to a decision variable *d*. This rule represents the cognitive strategy implemented by the model. Thereafter, the decision variable is mapped to a response *R* by adding Gaussian noise to its log-odds representation (Zhang & Maloney, 2012). The standard deviation of this noise distribution, denoted *σ*, was a free parameter in all models. The models differed only with respect to the strategy of the integration rule – the decision noise stage was identical in all of them.

**Figure 2.**
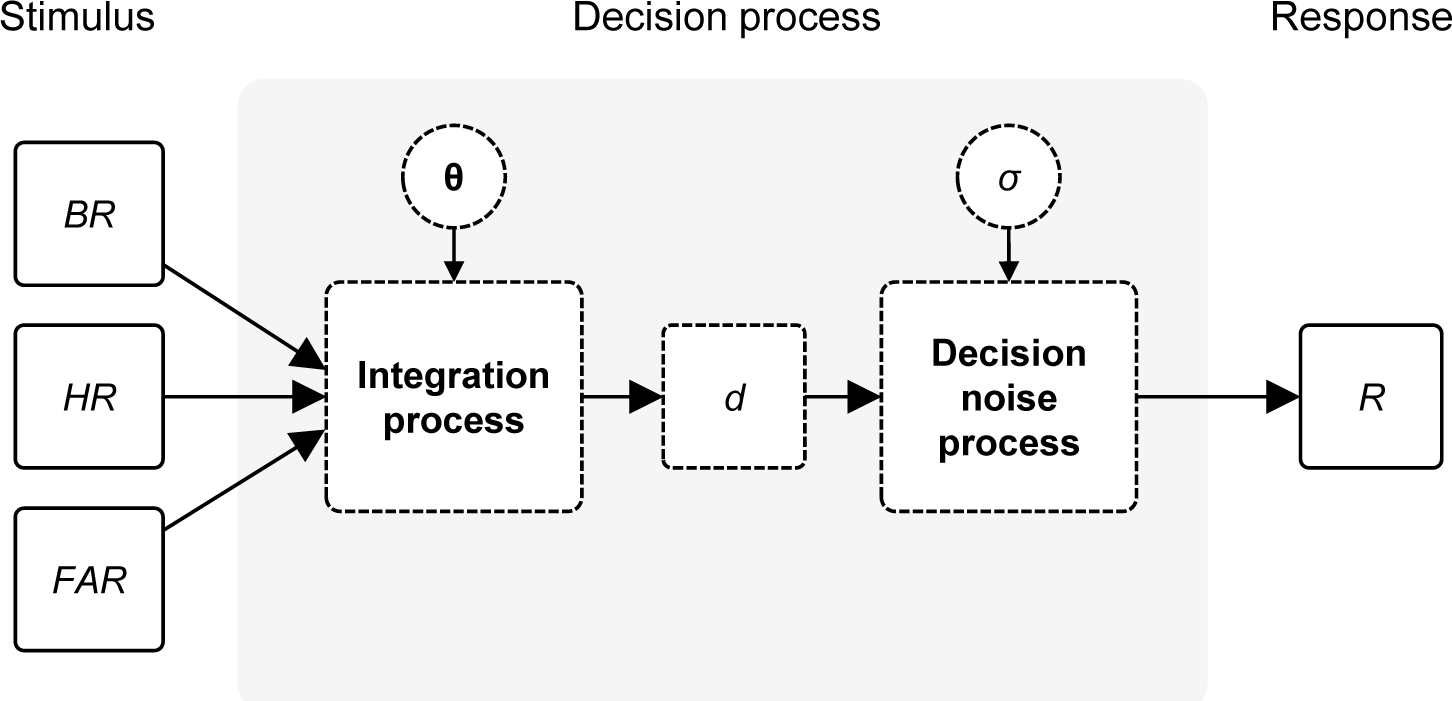
Schematic overview of the process models considered in this study. On each trial, the model receives three inputs (*BR*, *HR*, *FAR*). These inputs are mapped to a decision variable, *d*, through a deterministic integration process that differs between models. Finally, the decision variable is mapped to a response, *R*, by corrupting it with Gaussian noise. Vector **θ** specifies the model parameters related to the integration process and *σ* is the standard deviation of the late noise distribution. Note that in the tasks with the pictorial input format, we assume that there is also noise on the stimulus inputs before they are integrated (see text for details).

#### 2.5.1 Model 1: Bayesian integration with a prior

This model originates in a long tradition of normative theories arguing that human cognition is based on Bayesian inference strategies (Griffiths & Tenenbaum, 2006; Oaksford & Chater, 1994; Tenenbaum et al., 2011). The Bayesian strategy for the experimental task can be formulated in multiple ways. One way is to assume that the Bayesian observer simply applies Bayes’ rule to the provided BR, HR, and FAR information. The response of this model is

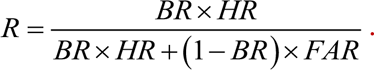

Even though this model uses the Bayesian judgement strategy, one could argue that it is non-Bayesian in the sense that it ignores any prior information it may have about base rates, hit rates, and false alarm rates of viruses and medical tests. Since we cannot rule out that participants have such priors, a proper Bayesian model should take this information into account. However, incorporating a prior for each of the three variables would introduce a relatively large number of free parameters to the model and make it overly flexible with respect to the other models. Therefore, we tested a simplified version in which the observer has a prior directly on the target probability (“the probability of having the virus given a positive test outcome”) rather than on the three input cues (BR, HR, FAR). We model this prior as a Beta distribution with parameters *a* and *b*, representing the number of previously observed number of positively tested people that do and do not carry the virus, respectively. Without additional information, this observer’s best estimate of the target probability is *a*/(*a*+*b*). To illustrate how this model updates its estimate after being presented with new information, consider the example in Figure 1B: 70 out of 100 people have the virus; 35 of the 70 persons with the virus got a positive test result; 9 of the 30 persons without the virus also got a positive test result. The observer would now have seen another *V*=35 cases of positively tested people who carry the virus and *X*=9 positively tested people who do not carry the virus. The new estimate of the target probability then is

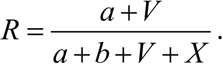

In the tasks where information is presented in the normalized format, we obtained *V* and *X* by transforming the stated proportions to frequencies under the assumption that they represent a sample of 100 cases. Although this may seem somewhat arbitrary, we note that increasing or decreasing the assumed sample size has a similar effect as decreasing or increasing, respectively, the values of *a* and *b*, which are fitted as free parameters. Hence, fixing the assumed sample size to an arbitrary value (rather than fitting it) is not expected to greatly affect the model fits, as long as we do not choose an extremely small or large value. Note that while this model captures the generic formulation of Bayes’ theorem stated above as a special case (at *a*=*b*=0), it also allows responses that deviate from this equation to be considered normative, in the sense that they take the uncertainty of the stated values into account.

#### 2.5.2 Model 2: Linear additive integration

The second model is based on findings from research on multiple-cue judgment tasks, such as estimating the price of an apartment based on its size, number of rooms, and distance to the city center (Brehmer, 1994; Juslin et al., 2008). This research suggests that people tend to combine cues by linearly weighting and adding them. Since the task in the present study can be seen as such a task – with *BR*, *HR*, and *FAR* as cues – it is conceivable that participants used this strategy. Under a linear-additive integration rule, the process model takes the form

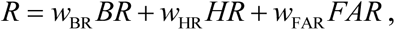

where the weights *w*_BR_, *w*_HR_ and *w*_FAR_ determine how much each piece of information contributes to the estimate of the posterior. The weights are fitted as free parameters with an unconstrained range.

#### 2.5.3 Model 3: An adapted heuristic toolbox

The third model is based on four heuristics that Gigerenzer and Hoffrage (Gigerenzer & Hoffrage, 1995) derived from self-reported strategies of a large number of participants performing a task similar to the one we use here. The first of these is the “joint occurrence” heuristic, which approximates the posterior as the product of the base rate and the hit rate, *d*=*BR*×*HR*. This will generally underestimate the true posterior, but can serve as a decent approximation when BR is high, which according to Gigerenzer and Hoffrage is the kind of situation in which people use this heuristic. The second heuristic entirely ignores *BR* and *FAR* and simply approximates the posterior as the *HR*, *d*=*HR*. This “Fisherian” heuristic leads to the same result as Bayes’ theorem when the base rate of the virus is equal to the base rate of positive test outcomes. Therefore, Gigerenzer and Hoffrage argue that people use this heuristic more frequently when the difference between these rates is small. The final two heuristics – referred to as “Likelihood subtraction” – are variants of an algorithm that ignores the base rate and seems to have been the predominant choice by participants in previous studies (Cosmides & Tooby, 1996). The first variant takes the difference between the hit rate and the false alarm rate, *d*=*HR*−*FAR*. The second variant is a simplification of this rule, in which the hit rate is assumed to be equal to 1, such that *d*=1−*FAR*. A challenge when implementing Heuristic Toolbox Theory is that there is no theory or proposed mechanism for how participants select the heuristic in each specific situation. This is a well-known problem, often referred to as the *strategy selection problem*. In related domains it has been suggested that people learn when to use what heuristic by feedback and reinforcement learning at the strategy level (Rieskamp & Otto, 2006), or that the ecological niches sometimes uniquely identify what heuristic that is applicable (Marewski & Schooler, 2011). Since this non-trivial problem has not yet been resolved for the base rate heuristics suggested by Gigerenzer and Hoffrage (1995), we decided to side-step it and focus on the more basic question of whether there is support for people using these heuristics, when they are evaluated in comparison with the other models. We do this in three ways: **i**) we consider a model assuming that people have learned to select for each situation the heuristic that best approximates the output of Bayes’ theorem (as inspired by Rieskamp & Otto, 2006). **ii**) In order to relax this ideal assumption of “well-adapted selection of heuristics”, we also consider a lexicographic version of the model that is more in line with the information integration emphasized in the Adaptive Toolbox program (Gigerenzer & Todd, 1999), and where the selection is based on how informative each cue appears in each situation (more in the spirit of Marewski & Schooler, 2011). **iii**) In the data analysis below, we finally also consider the most liberal application of the Heuristic Toolbox model that is possible, a disjunctive model, by allowing each response post hoc to be any of the five heuristics.

The model assuming that people have (somehow, to a decent approximation) learned to select the heuristic that best coincides with the output of Bayes’ theorem in each situation is,

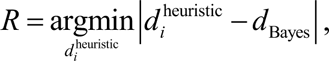

Where, 1 ≤ *i* ≤ 4, refers to the decision variables related to the four heuristic rules and *d*_Bayes_ is the optimal (correct) decision variable and |·| is the absolute value operator. The argmin operator selects the heuristic that most closely approximates the output of Bayes’ theorem.

#### 2.5.4 Model 4: A lexicographic heuristic toolbox

The final model that we tested is a “lexicographic” variant of the Heuristics Toolbox model (see Brandstätter et al., 2006; Gigerenzer & Goldstein, 1996, for models in this spirit). It uses the same four heuristics, but instead of choosing the heuristic that best approximates Bayes’ theorem in the given situation (Model 3), the lexicographic model assumes that participants consider the informativeness of each cue (*HR*, *BR*, *FAR*), in turn, until a sufficiently informative cue is found. The model takes an input probability as informative about the event’s occurrence when it deviates from 0.50 by more than a criterion value *δ*, which is fitted separately for *HR*, *BR*, and *FAR*. The order in which the model considers the cues is informed by the general pattern in the literature, which has shown that people respond strongly to hit rates, to some extent to the base rates, but rarely to the false-alarm rates (Juslin et al., 2011)^2^. Therefore, the model first checks the value of *HR*. If it is considered to be informative (i.e., deviates by more than *δ*_HR_ from 0.50), the participant reports *HR*, in accordance with the Fisherian heuristic introduced above. Otherwise, it next considers whether *BR* is informative about whether the event will happen and reports it if it is. If neither *HR* nor *BR* is sufficiently informative by itself and also the *FAR* is uninformative, the participant reports *HR*×*BR*, as according to the Joint Occurrence heuristic introduced above. Finally, if, on the other hand, *FAR* is informative, the participant reports *HR* – *FAR*, as according to the Likelihood Subtraction heuristic above. Formally, this model is formulated as

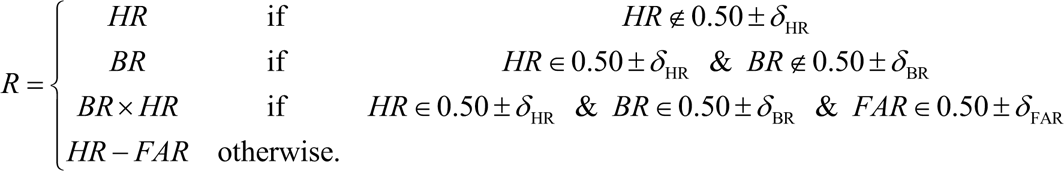

In this formulation, the informativeness of each cue is a step function of the possible cue-values: it is 0 when it deviates by less than *δ* from 0.50 and 1 otherwise. As a final step, we replace the step function with a sigmoid function, such that each cue has a certain *probability* of being considered informative. This probabilistic variant adds another parameter, *σ*_sigmoid_, that determines how quickly the probability of considering a cue as informative increases with its deviation from 0.50 (*σ*_sigmoid_ = 0 gives the original step-function formulation). This probabilistic formulation provides some additional flexibility to the model and can be interpreted as if participants have trial-to-trial noise in their informativeness criteria or do not apply them in a fully consistent manner.

#### 2.5.6 Decision noise

We added “late noise” to all models to account for various imperfections in neural information processing. We implemented this noise as a zero-mean Gaussian random variable with a free variance parameter, *σ*^2^. The noise was drawn independently on each trial and added to the log odds ratio of the model prediction on that trial,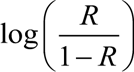, which was then transformed back to a probability^3^. Since the noise is implemented identically in all models, differences in goodness-of-fit are expected to reflect differences that models make in their assumptions about the judgment process.

Note that thanks to this noise, the “Probability theory plus noise” model suggested in earlier studies (Costello & Watts, 2014, 2016, 2017, 2018, 2019) is a special case of the Bayesian model we test here, namely the case of having no prior information (*a*=*b*=0). A novelty of the present work is that we allow the same kind of noise to corrupt judgments in the other models (“Linear-Additive Integration plus Noise”, “Heuristic Toolbox plus Noise”, and “Lexicographic Heuristic plus Noise”). This way, we can quantify evidence for the judgement mechanisms independently of the noise assumption.

#### 2.5.7 Parameter fitting and model comparison

We fitted the model parameters by using maximum-likelihood estimation, that is, by finding for each model the parameter vector **θ** that maximized 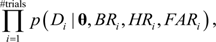, where *Di*, *BRi*, *HRi*, *FARi*, are the participant’s response and the presented base rate, hit rate, and false alarm rate on trial *i*, respectively. This maximization was done numerically, by using an optimization algorithm based on Bayesian direct search (Acerbi & Ma, 2017). Likelihoods were computed using Inverse Binomial Sampling (IBS) (van Opheusden et al., 2020). This is a numerical method that samples responses from the model until it matches the participant’s response; likelihoods are computed from the expected number of required samples until a match is found. An advantage of this method is that it guarantees the likelihood estimates to be unbiased. However, a disadvantage is that it can get stuck on parameter combinations that are unable to reproduce one or more of the participant’s responses in a reasonable amount of time. To avoid this, we added a freely fitted lapse rate *λ* to each model.

To avoid overfitting and account for differences in flexibility between models during model comparison, we fitted them using five-fold cross-validation. In each of the five runs, a unique subset of 20% of the trials was left out during parameter fitting. The log likelihood values of the left-out trials were summed across the five runs, providing a single “cross-validated log likelihood” value per model fit. For model comparison we computed the relative goodness-of-fit for each model as the difference in log likelihood with the best-fitting model. Since the optimizer may sometimes return a local maximum instead of a global maximum, we performed each fit 20 times with different starting points. We verified with a model recovery analysis that the models are distinguishable (see Figure A1 in Appendix).

#### 2.5.8 Estimating noise levels in the pictorial conditions

In the pictorial conditions, participants estimated the base rate, hit rate, and false alarm rate from the presented “probability matrices” (Figs. 1C-D). Previous research suggests that the amount of noise on these estimates scales with the numerosity of the estimated set (Piazza et al., 2004; Pica et al., 2004), which can be modelled using Weber’s law (Shepard et al., 1975). Instead of modelling this noise as an additional free parameter in the models for the pictorial conditions, we estimated it using an independently performed discrimination task (see *Procedure* above) and fitting a model that assumed that people’s observations of numerosity are corrupted by Gaussian noise that scales with the magnitude of the numerosity.

## 3. Results

### 3.1 Effect of base rate on responses

#### 3.1.1 Control experiment

The control experiment consisted of a single trial in which participants were presented with the classic formulation of the base-rate neglect task (base rate: 0.1%; hit rate: 100%; false alarm rate: 5%; correct answer: 1.96%). When counting all answers between 1.8% and 2.2% as correct (Sloman et al., 2003), 9% of the participants were classified as giving the correct answer (Figure 3A). This is consistent with a meta-analysis of 115 previously reported experiments, where the majority of the observed proportions of correct answers on his task was below 20% (McDowell & Jacobs, 2017). The modal response in our control task was “95%” (∼20% of the responses), which is also consistent with previous findings and which is often interpreted as base-rate neglect (Sloman et al., 2003). Hence, the control experiment successfully replicated earlier findings of base rate neglect. It is important to note, however, that while the *modal* response was 95%, the majority of the participants gave a correct or near-correct answer and were, thus, not neglecting the base rate.

**Figure 3.**
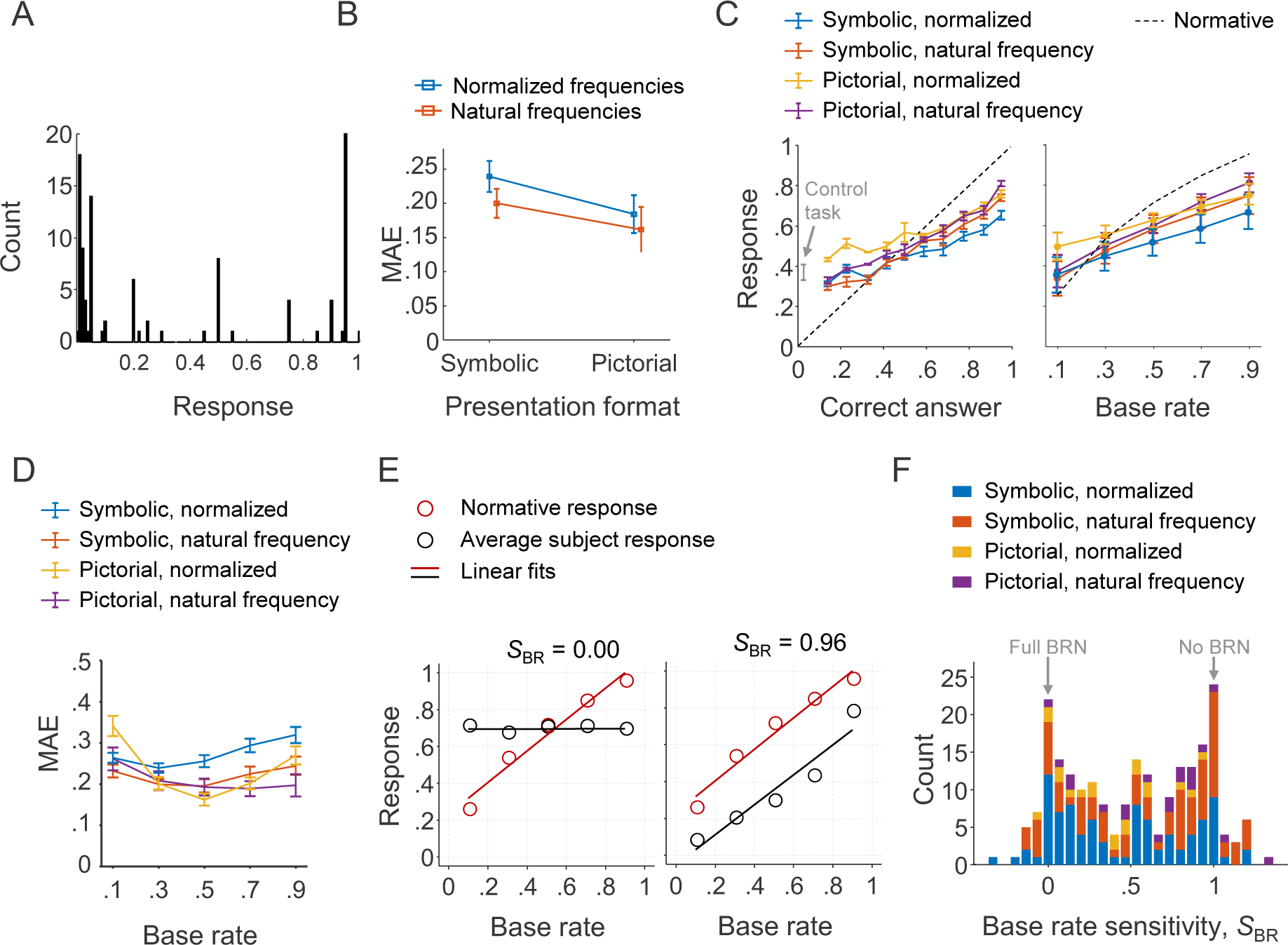
Analysis of participants’ sensitivity to base rates. (A) Distribution of responses in the control experiment. (B) Subject-averaged response accuracy in the main experiment, split by condition. MAE = Mean Absolute Error. (C) Left: Subject-averaged responses binned by the correct response and split by condition. Right: Subject-averaged responses for each base rate, collapsed across hit rates and false alarm rates and split by condition. (D) Subject-averaged response accuracy for each base rate, collapsed across hit rates and false alarm rates and split by condition. (E) Left: an example of a participant whose average response was independent of the base rate; the base rate sensitivity, *S*_BR_, was computed as the ratio between the linear fit slopes. Right: an example of a participant who responded to the base rate almost equally strongly as the normative observer. (F) Distribution of sensitivity values across all subjects. BRN = Base Rate Neglect.

#### 3.1.2 Main experiment

The task in the main experiment was the same as in the control experiment, except that it consisted of multiple items that varied in the presented base rates, hit rates, and false alarm rates. Since participants performed multiple trials, it is possible that they adjusted their responses based on their previous trials, which may have led to patterns in the data that might not have been there if they had performed only one trial. However, we found no evidence for such carry-over effects (Figure A2 in Appendix).

The response distributions (Figure A3 in Appendix) were similar to that of the single-item control experiment (Figure 3A), in the sense that for each item there was a large spread in responses but also a clear cluster of correct or near-correct responses.

Accuracy levels differed slightly between the four conditions (Figure 3B-C). Bayesian *t*- tests (JASP Team, 2020) confirmed that the participants’ average accuracy was reliably higher than that of a participant making a random guess on every trial (BF_10_ > 1.9·10^17^ in all four conditions)^4^. A two-way Bayesian ANOVA with mean absolute error as the dependent variable and task format and presentation format as independent variables revealed strong evidence for an effect of task format (BF_inclusion_ = 23.2), consistent with previous reports (Gigerenzer & Hoffrage, 1995) that people are more accurate when the information is presented as natural frequencies (*M* = 18.0, *SD* = 12.3) compared to normalized formats (*M =* 23.2, *SD* = 10.2). Moreover, the same test showed anecdotal evidence against both an effect of presentation format (BF_inclusion_ = 0.46) and an interaction effect (BF_inclusion_ = 0.37). Hence, somewhat unexpectedly, performance was comparable between the pictorial and symbolic conditions and the advantage of presenting the information as natural frequencies was also comparable between these two conditions.

If participants used a judgment strategy that ignored the base rate, then we should find that their average responses are similar across sets of trials with the same hit- and false-alarm rate values. Our data show that this is clearly not the case (Figure 3C right). In all four conditions, participants on average increased their responses in reaction to an increase in the base rate and hit rate and decreased their response in reaction to an increase in the false alarm rate (three-way repeated measures ANOVA^5^; BF_inclusion_ > 1000 for all main effects). Nevertheless, they did not adjust their responses as much as would be predicted from a normative perspective (dashed line in Figure 3C).

To examine base rate sensitivity at the level of single participants, we computed a sensitivity index, denoted *S*_BR_, as the slope of the linear fit to their responses divided by the slope of the linear fit to the responses from a Bayesian observer without a prior. A participant who entirely ignores the base rate has a sensitivity of 0, while a participant who reacts equally strongly to changes in the base rate as the Bayesian observer has a sensitivity of 1 (Figure 3E)^6^. We found that the participants’ sensitivity values are largely clustered around 0 and 1 (Figure 3F), which suggests that many of them either fully ignored or fully accounted for the base rate. This finding is qualitatively consistent with the data obtained using the classic base-rate neglect task, where we found a similar kind of clustering (Figure 3A). In addition, we found that the base-rate sensitivity index correlates with median log response times (*r* = 0.20, *p* = 0.004). Such a correlation is expected when considering that heuristics are supposed to be “fast and frugal” (Gigerenzer & Todd, 1999) but also ignore part of the provided information.

Finally, we noticed that some of the participants had extremely short or extremely long median response times (range: 3.08 to 69.1 seconds). We suspect that extremely fast participants might have been “clicking through” the task and extremely slow ones might have been using external resources. To verify that our results do not critically depend on data from such participants, we redid the analyses after filtering out participants with extreme median reaction times and found that the results are similar and lead to the same conclusions (see Figure A4 in Appendix).

We draw two conclusions from the results so far. First, there are considerable individual differences in how participants behave in both the classic formulation of the task and our generalization of it: in both tasks, there is a group of participants that almost entirely seems to account for base rates in their judgments and another group that almost entirely neglects them. This heterogeneity in behavior warrants caution when attempting to make population-level statements about base rate neglect. Second, the base-rate neglect effect does not seem to be limited to inference problems with a high hit rate and extremely low base rate.

### 3.2 Model comparison

To get more insight into the kind of judgement strategies that the participants may have been using, we fitted four models to their individual data sets: a Bayesian model, a linear-additive integration model, and two heuristic models. These models represent distinct theories about cognitive strategies and make clearly distinguishable predictions (Figure A1 in Appendix). A key difference is that the former two models integrate the cues while latter tend to simply report one of the cues rather than integrate them. While it would be overambitious to expect that fitting 4 models to 48 responses can reveal the exact strategy used by each participant, it can still provide useful insights into the *kind* of strategies they were using. For example, a participant who integrated the three cues is expected to be captured well by the Linear-Additive model and, possibly, also by the Bayesian model (if the integration rule is close to the Bayesian one), but not by the heuristic models. Likewise, participants using a non-integration strategy are expected to be captured very poorly by the Bayesian model and, possibly, well by the heuristic models.

Most participants in the experiment with symbolic stimuli were best captured by the Bayesian and Linear-Additive models, a few of them were best captured by the Lexicographic Heuristic, and almost no participant was described well by the Heuristic Toolbox model (Figures 4A and 5A). Moreover, all models did a good job at capturing the participants’ group-averaged responses as a function of the base rate (Figures 4B and 5B). However, consistent with the model comparison results, the Linear-Additive and Bayesian models clearly outperformed the heuristic models when considering fits at a much more fine-grained level (Figures 4C and 5C). These results thus suggest that most participants integrated the cues, with some participants apparently using an integration rule that resembled the Bayesian one.

**Figure 4.**
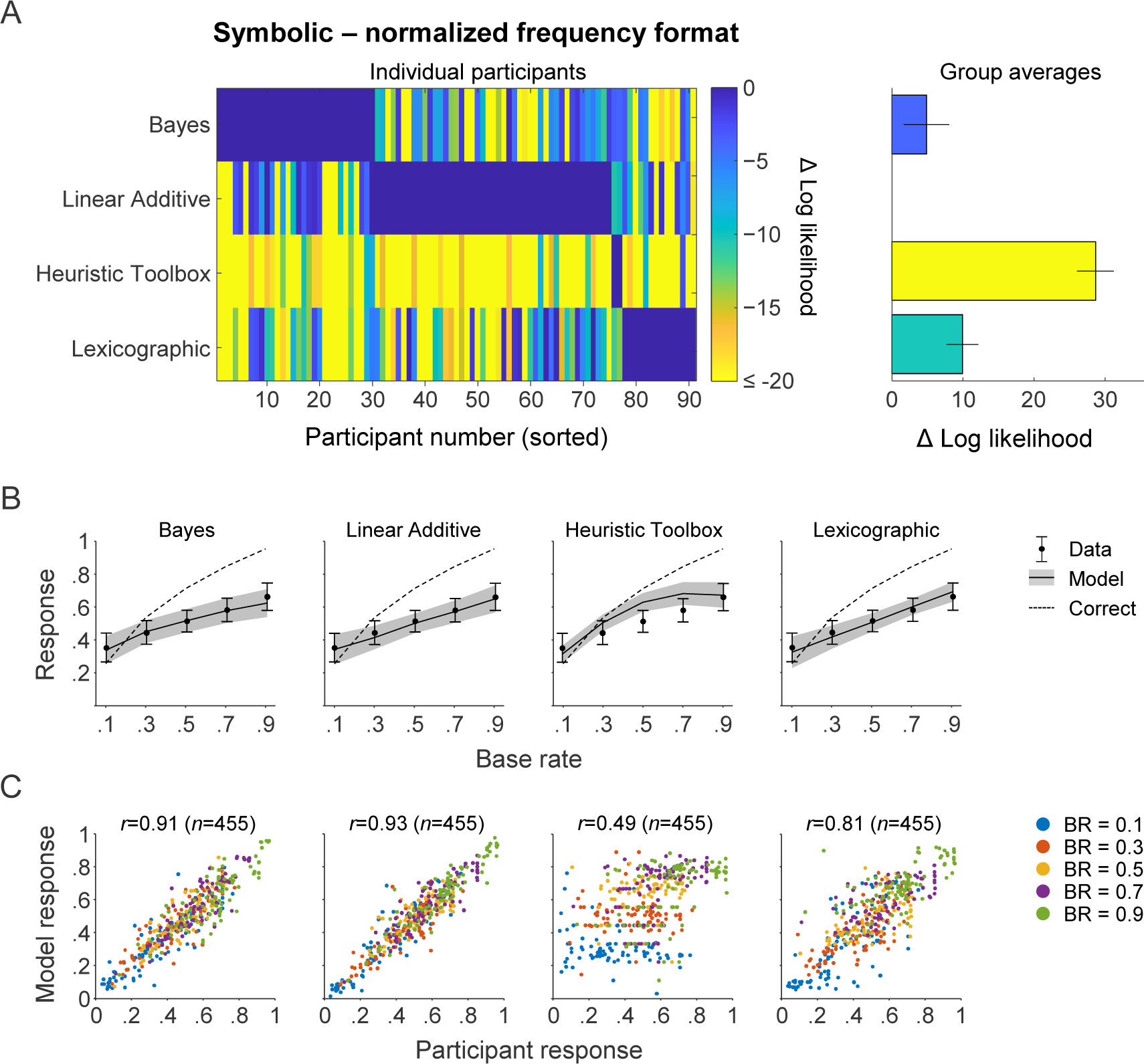
Modeling results for the condition with symbolic stimuli presented in normalized format. (A) Left: Cross-validated model log likelihoods relative to the best model. The preferred model for each participant is indicated in blue and the worst models are indicated in yellow. For visualization purposes, participants were sorted in such a way that all participants on which a particular model was the preferred one would line up (blue areas). Right: Cross-validated model log likelihoods relative to the Skeptic Bayesian model, averaged across all participants. Errors bars indicate 1 s.e.m. (B) Participant-averaged responses as a function of the base rate. Error and shaded areas indicate 1 s.e.m. (C) Simulated model responses (including late noise) under maximum-likelihood parameters plotted against participants responses. Each point represents the average response across the 9 trials within a base rate (color-coded); *r* indicates the Pearson correlation coefficient and *n* the number of points in each plot. The model order is the same as in panel B.

**Figure 5.**
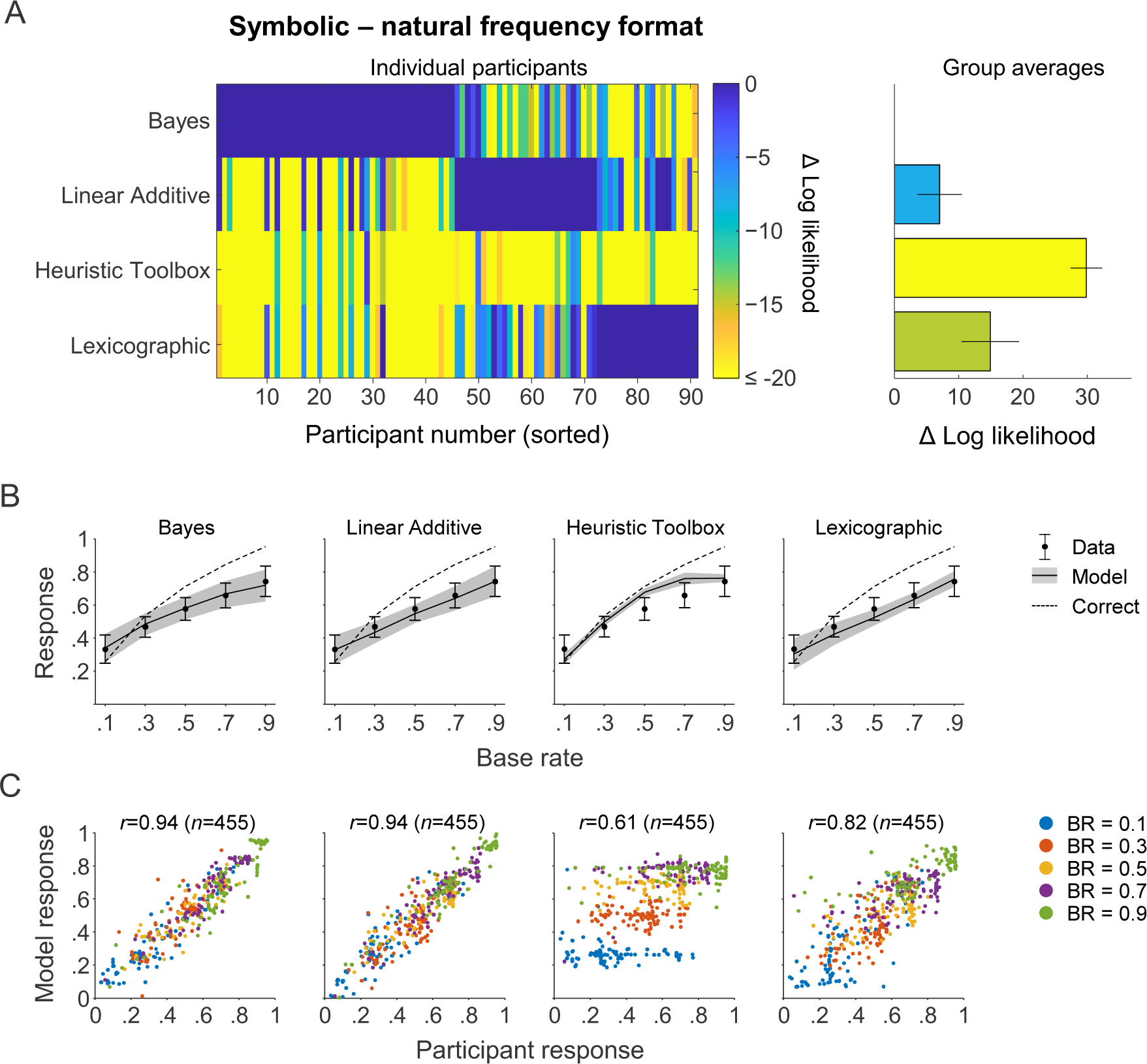
Modeling results for the condition with symbolic stimuli presented in natural frequency format. This figure follows the same layout as Figure 5.

Just as in the base-rate sensitivity analysis above, we found large individual differences in model preference. Part of this variability may be due to model evidence itself being a random variable, in the sense that measuring the same participant twice on the exact same task will not result in identical log likelihood values. However, for many participants one model was preferred strongly over the other three models, which suggests that at least part of the variability in model evidence is due to different participants using different strategies. While not of primary interest in the present study, it is interesting to note that more participants were classified as a Bayesian in the natural frequency format (46 out of 91) compared to the normalized frequency format (31 out of 91), *p*=0.035 (Fisher’s exact test). This is consistent with earlier literature as well as with the accuracy difference reported above (Figure 3B).

The modeling results in the pictorial conditions were very similar to those in the symbolic conditions. The Bayesian and Linear-Additive models again captured the data convincingly better than the heuristic models (Figures 6A and 7A) and provided good fits to the data (Figure 6B, 6C, 7B, 7C). Moreover, there were quite a few participants for whom the three non-winning models provided substantially worse accounts of the data than the winning model, which is suggestive of individual differences in strategies. However, we do not see a shift towards the Bayesian model in the task with the natural frequency format (11 out of 20) compared to the normalized frequency format (10 out of 20), *p* = 1.00 (Fisher’s exact test).

**Figure 6.**
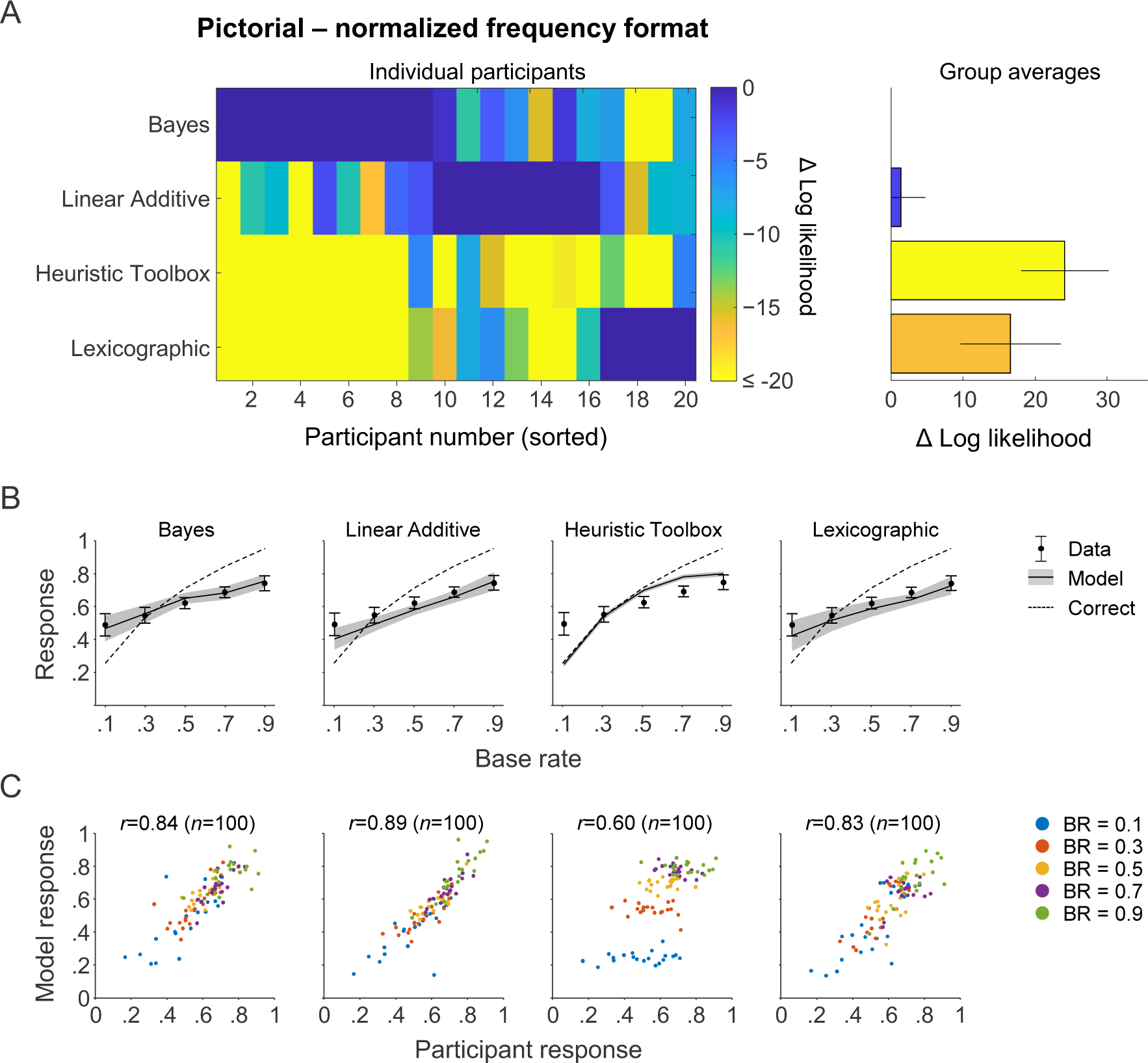
Modeling results for the condition with pictorial stimuli presented in normalized format. This figure follows the same layout as Figure 5.

**Figure 7.**
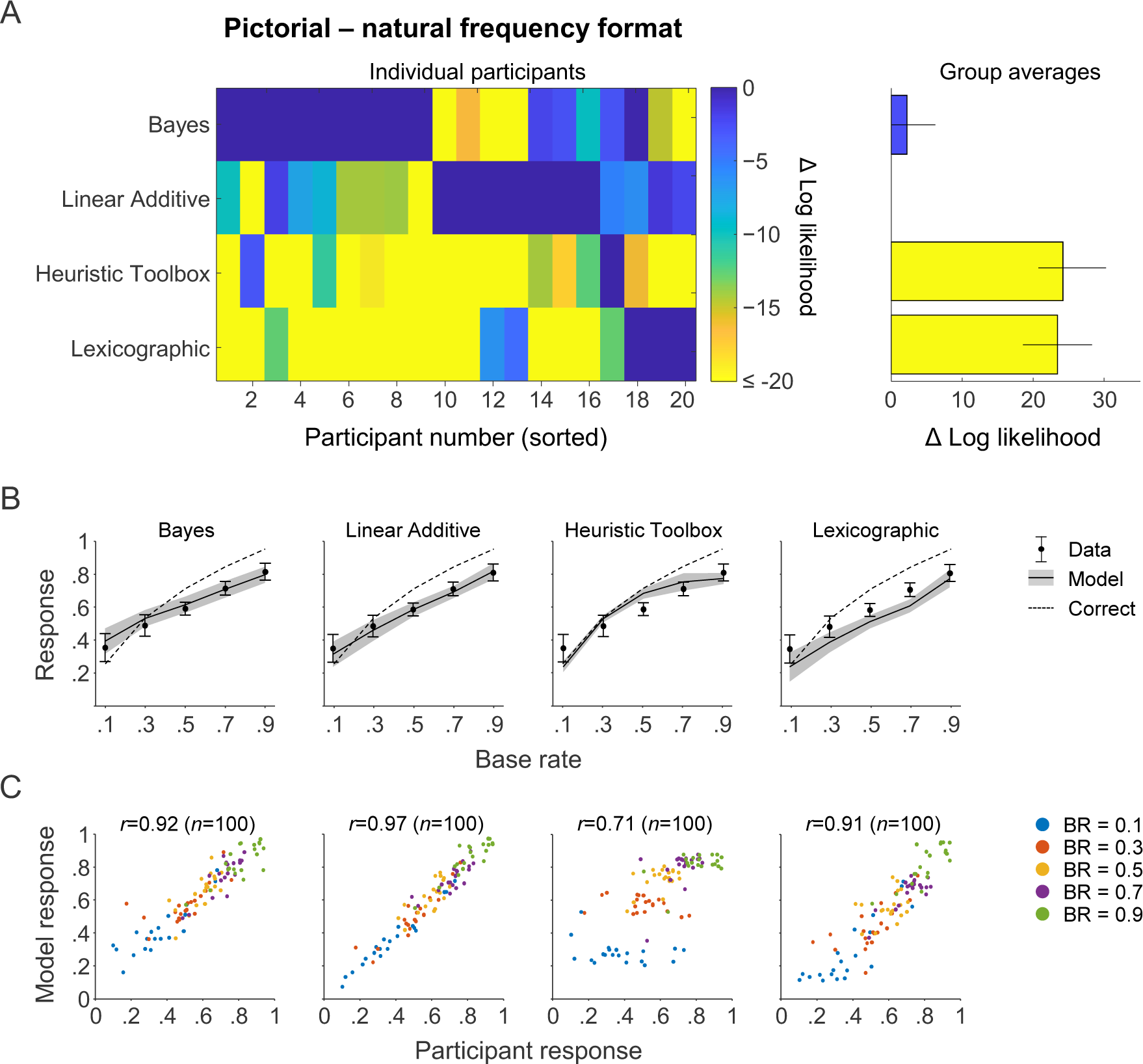
Modeling results for the condition with pictorial stimuli presented in natural frequency format. This figure follows the same layout as Figure 5.

To further verify that the Linear-Additive model and Bayesian model represent mathematically distinctive decision strategies, we examined the correlation between their predictions under maximum-likelihood parameter estimates. The results (Figure A5 in Appendix) show that the predicted responses are highly correlated, which was expected, because both models were fitted to follow the empirical responses. Importantly, however, on many trials the models make quite different predictions, which verifies that they implement mathematically distinct strategies^7^. This is consistent with the observation of many participants with quite distinct differences in model fit between the two models in Figures 4 to 7.

Finally, we verified that the modeling results do not critically depend on the chosen noise distribution, by rerunning it using a different noise distribution. This gave very similar results (see Figure A6 in Appendix).

The modeling results support our earlier conclusion that there are individual differences in the strategies employed by the participants. Moreover, they provide very little evidence for heuristic judgment strategies.

### 3.3 Analysis of the relation between judgement strategies and base rate sensitivity

The results so far suggest that there are individual differences both in the extent to which participants accounted for base rates in their judgments (Figure 3F) and in the preferred cognitive model fitted to their data. We next investigated whether these two findings may be related: is there any evidence in the modeling results to suggest that participants with little base rate neglect were using a categorically different strategy than participants with strong base rate neglect? To examine this, we replotted the distribution of sensitivity values shown in Figure 3F, but now color-coded by the best-fitting model for each participant (Figure 8). While both models span a broad range of sensitivity values, we found that the Bayesian model was preferred mainly for participants with little or no base-rate neglect (i.e., *S*_BR_ values close to 1), while the Linear-Additive model was more successful in capturing data from participants with strong base-rate neglect (i.e., *S*_BR_ values close to 0). This finding suggests that participants with little base-rate neglect may have been using a categorically different reasoning strategy than those with strong base-rate neglect.

**Figure 8.**
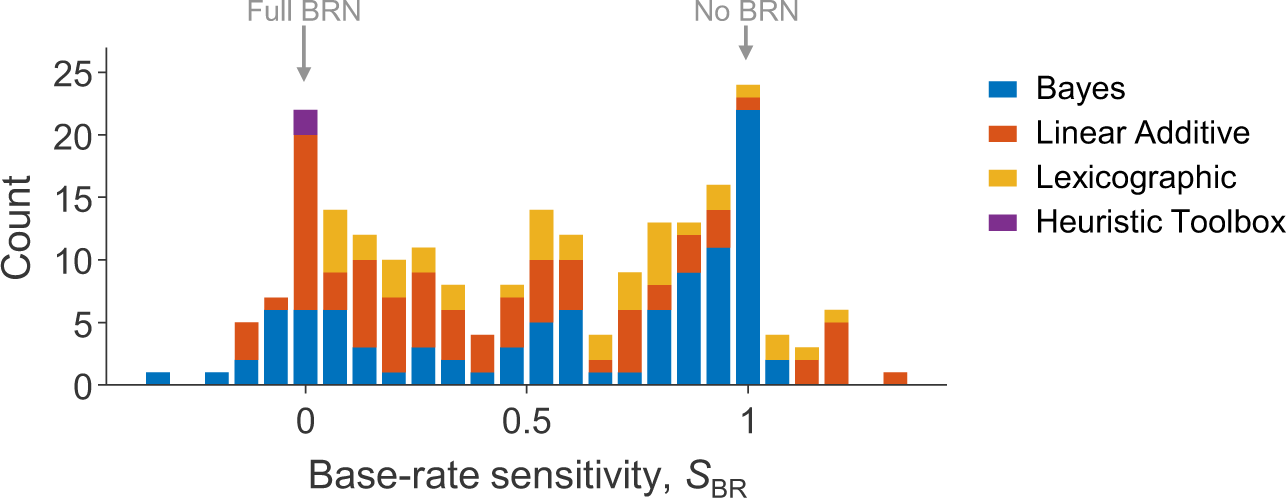
Distribution of base-rate sensitivity split by best-fitting model. Participants who showed little or no base-rate neglect are generally best accounted for by the Bayesian model, while participants with strong base-rate negelect are generally better accounted for by the Linear-Additive model.

**Figure 9.**
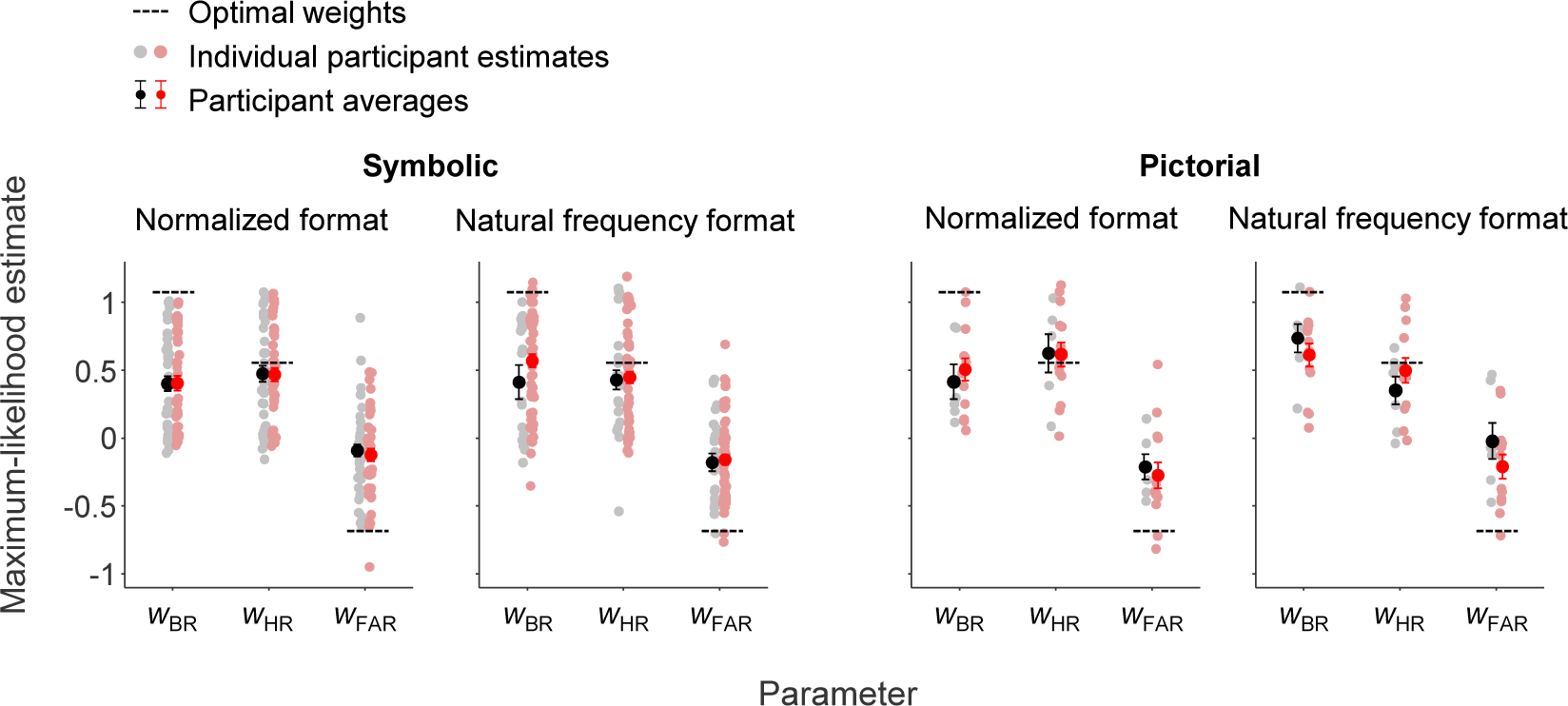
Estimated weights of the linear additive model. Estimated weights for the base rate, hit rate and false-alarm rate for the linear additive model. Black error bars and dots indicate the mean and individual weights of participants for whom the linear additive model provided the best fit. Red error bars and dots indicate the mean and individual weights of all other participants. The dashed lines indicate the weights of an additive integration model fitted to responses from a Bayesian decision maker (i.e., the linear additive weights that best approximates the Bayesian solution).

### 3.4 Analysis of model parameter estimates

To further evaluate the model fits, we next looked at the estimated parameter values. We limited this analysis to the key parameters in the Bayesian and the Linear-Additive integration models, which together accounted for 80% of the participants in terms of the best-fitting model.

*Weights in the Linear-Additive model.* According to the fits of the Linear-Additive model, participants gave on average too little weight to both the base rate and false-alarm rate in all four conditions when compared to the optimal weights (i.e., the weights that would have minimized the RMSE in our task). In particular the false-alarm rate was underweighted, which is a finding that can be linked to several other robust phenomena in psychology, such as pseudo- diagnosticity (Ofir, 1988) and subadditivity (Tversky & Koehler, 1994). The hit rate, on the other hand, was on average weighted close to optimally.

*Prior in the Bayesian model*. In the Bayesian model, the estimated median number of previously observed cases (i.e., the median of the sum of the *a* and *b* parameters) was 62.6 (*Q*_1_=9.30; *Q*_3_=335) when considering all participants and 10.2 (*Q*_1_=0.51; *Q*_3_=422) when considering only the participants for whom the Bayesian model was the best-fitting model (*Q*_1_ and *Q*_3_ refer to the first and third quartile). According to these results, participants either had a very weak prior or one of a strength corresponding to having observed a few hundred earlier cases. Simulation results (see Figure A7C in Appendix) show that priors in this range can produce regressive effects of similar magnitude as observed in the data (Figure 3C). Hence, a deterministic “dampening with a prior” seems a more plausible explanation for the regressive effects found in our data than a classical regression effect caused by random noise.

*Noise levels in both models.* The median of the estimated noise parameter *σ* was 0.37 (*IQR* = 0.54) for participants best fitted by the Bayesian model and 0.43 (*IQR* = 0.48) for those best fitted by the Linear-Additive model^8^. This may seem large, but we remind the reader that the noise was applied to the log-odds ratio of the predicted responses. For comparison, a noise value of 0.37 on the log-odds ratio corresponds to Gaussian noise with a standard deviation of 0.09 on the raw response. Simulation results (see Figure A7B Appendix) show that this amount of noise is far too small to cause regressive effects of the magnitude seen in the data (Figure 3C). Hence, decision noise does not seem to be a plausible explanation for the regressive effects found in our data, but they seem better accounted for by a dampening with a prior.

### 3.5 Frequency of heuristic judgments

The Heuristic Toolbox and Lexicographic models tested above incorporate a variety of heuristic judgement rules (“Report HR”, “Report 1 − FAR”, etc). We found that both models provide relatively poor descriptions of the data. One possible explanation is that participants did generally not use the tested heuristics in their judgements. However, another possibility is that participants used a selection rule (which heuristic to use on a given trial) that differed from the selection rules in the Heuristic Toolbox and Lexicographic models. To investigate this, we evaluated a post hoc disjunctive model and counted for each participant in the two conditions with symbolic stimuli how many of their responses coincided with the output of any out of five heuristics. Even though not used in any of the tested models, we added “Report FAR” as an additional heuristic in this analysis. The results (Figure 10) showed that in both conditions approximately half of the responses were consistent with the disjunction of the five heuristics, while the other half were inconsistent with all five heuristics. Hence, the best-fitting model of all possible models based on these five heuristics can explain at most half of the responses. Because several of the heuristics coincide with merely reporting one of the numbers stated in the problem it may also be hard to verify that all of these responses are deliberate and intentional inferences referring to the posterior probability, as opposed to more superficial response biases. Or, put differently, if participant responses are exclusively based on heuristic strategies, then there must exist other, yet to be discovered heuristics that were not included in the current analysis.

**Figure 10.**
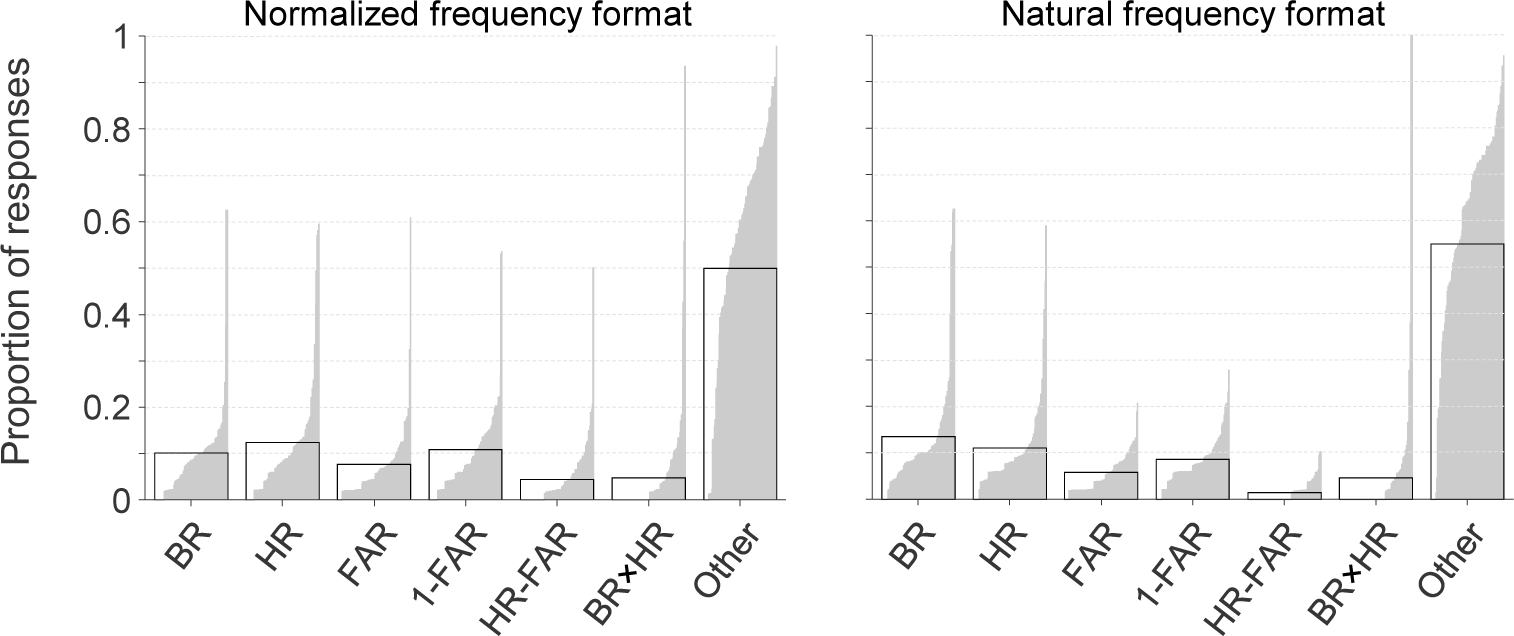
Distribution of participant responses in the two conditions with symbolic stimuli. The proportion of participant responses that were equal (in two decimal places, when expressed as a probability) to a response predicted by one of the heuristic rules. Each bar corresponds to one participant. Responses that corresponded with more than one heuristic were counted multiple times, which means that some of the counts may be slightly overestimated. The black rectangles indicate the mean for each group of responses. In both conditions, approximately half of the responses coincided with one of the heuristics.

## 4. Discussion

In this study, we examined the generality and cognitive basis of the base-rate neglect phenomenon in the context of the medical diagnosis task. To this end, we tested participants on the medical diagnosis task under a large range of different combinations of base rates, hit rates, and false alarm rates. Our empirical results indicated large individual differences in the degree with which participants neglected base rates. This heterogeneity was reflected in the cognitive modeling results, where the evidence was divided mainly between the Bayesian and Linear-Additive models and provided little evidence for heuristic strategies.

### 4.1 Generality of the base-rate neglect effect

We found that signs of base rate neglect were present throughout the tested space of problems and to a similar degree as found in the classical task. Hence, the phenomenon known as base-rate neglect does not seem to be limited to inference problems with extremely low base and false-alarm rates in combination with a high hit rate. Taken as a whole, participants on average seemed to make proper use of the hit rate but underused the base rate and false-alarm rate.

Interestingly, however, we found large individual differences in behavior. In particular, both in the control experiment (using the classical task) and in the main experiment (using the generalized task), we found two clusters of participants: those who seemed to largely ignore the base rate and those who seemed to account for it well. This heterogeneity suggests that there may be true individual differences in rational thought (Stanovich & West, 2000), which would warrant caution in drawing strong group-level conclusions about base rate neglect. Instead, the phenomenon may be best studied at the level of individuals.

Finally, even though the symbolic and pictorial tasks at first glance might seem rather different, the participants on average seemed to use the same kind of strategies to solve the tasks. This is an interesting finding since even though the underlying task structure was identical in the symbolic and the pictorial versions of the task, the judgments were based on exactly stated numbers in the symbolic condition and on uncertain assessments in the pictorial condition demanding the participants to reason under uncertainty.

### 4.2 Cognitive basis of the base-rate neglect effect

The heterogeneity in empirical estimates of the degree of base-rate neglect was reflected in the model comparison results. Even though we found little evidence for the heuristic models, evidence between the Linear-Additive and Bayesian models was divided: the data of some participants were better accounted for by the former model, while data for others were better accounted for by the latter.

More specifically, participants who showed little or no base-rate neglect were generally best described by the Bayesian model. The estimated prior counts in this model varied from near-zero to a few hundred cases, which suggests that people had weak to moderately strong priors, but rarely so strong that they completely dominated the data given in the experimental trials. Interestingly, the noise levels were estimated to be relatively small. Simulations revealed that much higher noise levels would have been required to explain the typical regressive effects found in empirical data. Hence, our results suggest that those effects were largely caused by “dampening” due to the prior, which is consistent with a proposal in other recent papers (e.g., Zhu et al., 2020).

Participants who showed strong base-rate neglect effects were generally best described by the linear-additive model. From the perspective of that model, participants have a qualitative understanding that both the evidence and base-rate is relevant in base-rate problems, but – as often observed in other multiple-cue judgments tasks (Juslin et al., 2008, 2011) – they spontaneously add up the cues, rather than engaging in the multiplication prescribed by probability theory (Juslin et al., 2009). This suggests that from their experience with the world, people may have obtained normative insights at a general qualitative level, but they are unable to perform the normative mathematical integration when presented with symbolic representations of probability. This interpretation is vindicated by the observation that even after explicit tutoring and instruction on how to compute the posterior probability from Bayes’ theorem in base-rate problems, people are still better described by linear additive integration models than by the normative integration model (Juslin et al., 2011). One potential criticism of the Linear-Additive model concerns its flexibility, in the sense that it can produce a wide array of apparently different strategies, ranging from exclusively weighting the hit rate or base rate, to integrating pairs of components, to integrating all three components in approximating to Bayes’ theorem. One could, however, argue that this captures the richness and variety of the strategies found in human participants and could reflect individual differences in, for example, their knowledge of probability theory and their judgment of the relevance of various contextual cues (Ajzen, 1977; Bar-Hillel, 1980; Birnbaum & Mellers, 1983; Fischhoff et al., 1979; Fishbein, 2015; Goodie & Fantino, 1999). The strategies are effectively identified by the parameters of the model and the common cognitive claim by the model is that people have difficulty with “number-crunching” symbolic representations of probability according to Bayes’ theorem, but rather often default to linear additive integration of the components.

Data from the remaining ∼20% were best captured by the heuristic models. Heuristics come in many varieties and the individual strategies (e.g., “multiply hit rate with base rate”) are not assumed to be used for all tasks but rather to be picked when the structure of the task at hand fits some criteria (e.g., “use the joint occurrence heuristic if the base rate is high” (Gigerenzer & Hoffrage, 1995). However, the exact conditions for when the different heuristics should be used are difficult to establish and have not been specified for the heuristics in these base-rate problems. For example, how high does the base rate has to be for it to be considered high? In this study we used two different heuristic models that circumvented this problem in different ways. While there may remain untested heuristic models with selection rules that differ from the ones we tested here, we could see from the distribution of responses (Figure 10) that *any* heuristic model using the strategies considered in the present study can account for at most approximately 50% of the responses. Hence, if judgments in the medical diagnosis task are mainly driven by heuristics, there must be additional, yet-to-be-discovered strategies that participants were using. A challenge for the heuristics program is to uncover these strategies as well as the mechanism that determines which strategy to use in which situation.

Another challenge is to find the source of the individual differences in the strategies. These individual differences may involve different choices of strategies (e.g., report of a single cue *vs* Bayesian integration of all cues) as well as differences in how a specific strategy is applied (e.g., what cue is selected or what priors are entered in the Bayesian integration). These differences may, in turn, derive from differences in how the participants represent the task and in what they consider to be important aspects in the task (Szollosi & Newell, 2020), as well as from individual differences in fundamental cognitive abilities, such as intelligence or working memory (Conway & Kovacs, 2013) or differences in how much experience participants had with this kind of task, with novices relying more on heuristics than experienced participants. In the present study, however, we are not in the position to further explore this important question.

### 4.3 Effect of frequency format and presentation format

In the symbolic conditions, using a natural-frequency format led to slightly better task accuracy. In addition, more participants in those conditions were classified as Bayesian when presented with information in the natural-frequency format (21 vs 11). Nevertheless, the differences were not as drastic as in some previous studies. The original study by Gigerenzer and Hoffrage found that changing from a normalized frequency format to a natural frequency format increased the performance rate from 16% correct responses to 46% correct responses (Gigerenzer & Hoffrage, 1995). A later study by Cosmides and Tooby found similar results, with a performance increase from 12% to 56% correct responses (Cosmides & Tooby, 1996). One explanation for why the increase in the present study was smaller is that our participants were required to give their answer in percentages rather than a ratio. An alternative explanation is that the task structure did not make the set relations between base rate, hit rate and false-alarm rate transparent enough (Barbey & Sloman, 2007; Sloman et al., 2003). The nested-sets hypothesis stem from the dual-process model and – in contrast to Gigerenzer and colleagues’ view that people are helped by the natural-frequency format in and of itself – the proponents of this hypothesis argue that the format is only beneficial because it can make the task structure more transparent. In this view people use two systems to reason, one primitive associative judgment system that sometimes leads to errors in judgment and a second more deliberate rule-based system. The use of the second system is only induced if the task is represented in a way that is compatible with the rules. In this case the rules are elementary set operations which means that the problem needs to be readily formulated in terms of sets. The problem becomes even easier if the relevant sets are all nested (Sloman et al., 2003). In other words, if the chance of having the disease is nested within the chance of testing positive which in turn is nested within the set of all possible cases. Since in our task the hit rate was unequal to 1 it means that the sets are not nested (the chance of having the disease is not a subset of having a positive test) and therefore the task structure was less beneficial for the participants.

### 4.4 Symbolic vs pictorial presentation

Even though the pictorial format introduced uncertainty in the task by forcing the participants to make their own estimations, this did not affect the overall performance. There is a trend towards better performance in the pictorial format (Figure 3), but all in all there was anecdotal evidence against an effect of presentation format. This also means that the pictorial format did not function as a visual aid benefiting the participants as some previous studies has found (unless these two effects happened to cancel each other out). However, previous studies on the effects of visual representations have received mixed results and in the cases where a pictorial representation has been beneficial it was only presented as an addition to the question already using symbolic formulations (Brase, 2009; Garcia-Retamero & Hoffrage, 2013) and never by itself as was the case in our task. We can only speculate about the reason that the two formats did not lead to large differences in performance. One intriguing possibility is that the mental representations of the symbolic numbers may have been just as uncertain as those of the pictorial estimates. It is also possible that there are multiple factors working against each other.

The task given in the symbolic format is similar to a typical math word problem and could therefore cause people with high levels of math anxiety to perform worse than they would have if they had received the pictorial task instead (Luttenberger et al., 2018).

### 4.5 Limitations and future directions

The results of the linear-additive model fits suggested that participants – on average – strongly underweighted false-alarm rates, possibly even more so than base rates. The fallacy of false-alarm-rate neglect has received support in previous studies (Ofir, 1988) and it would have been interesting to investigate in a more extensive stimulus space. The current design, however, was tailored to studying sensitivity to base rates and was less suitable for studying sensitivity to hit rates and false alarm rates, because of the more limited ranges for those two rates included in our experiments.

Since there are a number of factors that differ between data collected in a lab and data collected online, one has to be careful when comparing the results. Even though it is impossible to make sure that the data has been collected under similar conditions it is at least possible to compare the data after it is collected. In a separate data collection for a different study (not reported here), lab participants were presented with the exact same task as we presented to the online participants in the present study, which enables a direct comparison between the two data collection methods. A Bayesian independent t-test showed anecdotal evidence in favor of there being no difference in the mean absolute error between the two groups (BF_01_= 1.45), which suggests that performance levels are comparable between the two kinds of participants. Just as in previous work, we found a benefit of the natural frequency format on performance. This effect has been explained by either referencing how during the evolution of the human race, naturally sampled frequencies is what we have been exposed to and subsequently evolved to use, or one has attributed the beneficial properties to the clarification of the task and thus making it easier for people to use deliberate thought instead of associative decision making. These two accounts are focused on how the difference in format makes people switch from one strategy towards using another one. There is, however, also a possibility that the switch in format does not cause people to switch strategies but to use the same strategy with better tuned parameters. For example, a participant might shift from using a heuristic to a Bayesian strategy when the format changes from normalized frequencies to natural frequencies, or they might use a linear additive strategy in both formats but use better tuned weights. Our use of a between-subject design is a limitation in this respect. Future work could study the effect of frequency format using a cognitive-modelling approach applied to within-subject data. That would allow getting a more detailed insight into the nature of the reasoning strategies people use.

### 4.6 Conclusion

In this paper, we investigated the generality and cognitive strategies responsible for the phenomenon known as “base-rate neglect”. We found that the phenomenon generalizes to reasoning problems beyond those typically used in previous studies. However, we also found evidence for substantial individual differences in the degree of base-rate neglect. Our modelling results suggest that these individual differences may reflect individual differences in the cognitive strategies employed by the participants. An interesting direction for future research is to investigate this possibility in more detail.

## APPENDIX

### Model recovery analysis

We performed a model recovery analysis to examine how well the models can be distinguished from each other. In this analysis, we generated 20 synthetic datasets from each model by simulating responses to the same 45 trials as were shown to our human participants. To make synthetic datasets representative of the empirical data, we performed these simulations with parameter values set to the maximum likelihood estimates of randomly chosen participants. If two or more of the models make very similar predictions, then the recovery analysis will have difficulties in determining which model generated which dataset. The results (Figure A1) show that this is not the case: for each of the four groups of synthetic datasets, the model that generated the data is reliably selected as the one that best accounts for them. This indicates that the models make clearly different predictions and that our methods are adequate in detecting these differences.

**Figure A1.**
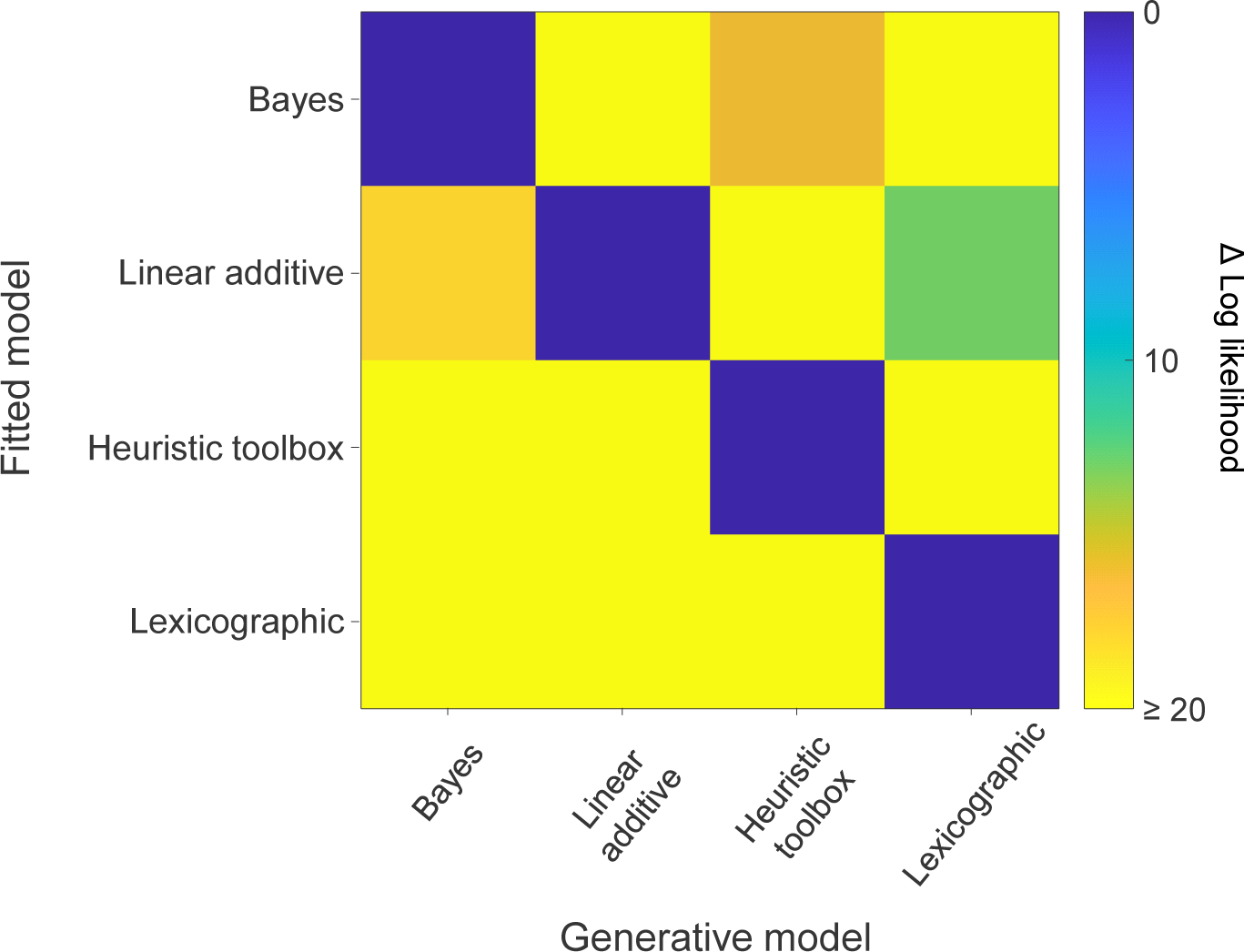
Model recovery. Average cross-validated log likelihood values across twenty generated datasets from each model, relative to the best-fitting model. To ensure that the generated datasets were representative for empirical data, they were generated with maximum-likelihood parameters from randomly selected human participants and consisted of the same 45 trials as presented to the human participants. The results show that the models make clearly distinct predictions: each model is the best-fitting model for its own data and provides a poor fit to the data from other models.

### Analysis of carry-over effects in the main experiment

Since the participants performed multiple trials, it is possible that they adjusted their responses based on their previous trials, which may have led to patterns in the data that might not had been there if they had performed only one trial. To examine whether this was the case we analyzed performance as a function of trial number in the MTurk data. These data had been collected in such a way that every item was presented as the first trial for at least two participants, as the second trial for two other participants etc. If performing multiple trials improved performance, the correlation between participants’ responses and the correct responses should increase over trials. This was not the case (Figure A2). We also compared the *RMSD* of the first trial with that of all consecutive trials. For both the normalized- and natural frequency format, a one sample Bayesian *t*-test showed that the *RMSD* of the first trial is not larger than the rest (BF_0-_ = 19.70 and BF_0-_ = 61.86 in favor of the null hypothesis). Both these results suggest that the judgement strategies employed by the participants were stable over trials, including the first one.

**Figure A2.**
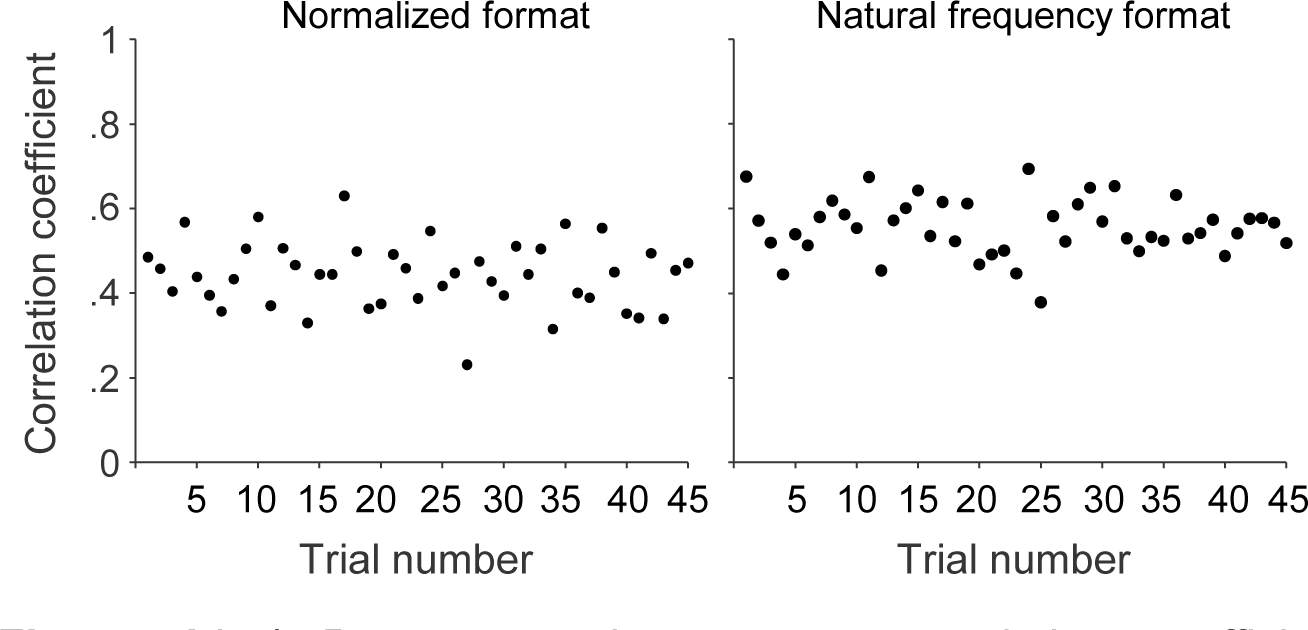
Pearson product-moment correlation coefficients between the participants’ responses and the correct responses, as a function of trial number. Correlations were computed separately for every trial number. Left: Normalized format. Right: Natural frequency format.

### Response distributions in the main experiment

Figure A3 presents the response distributions in the main experiments, separately for each of the 45 items.

**Figure A3.**
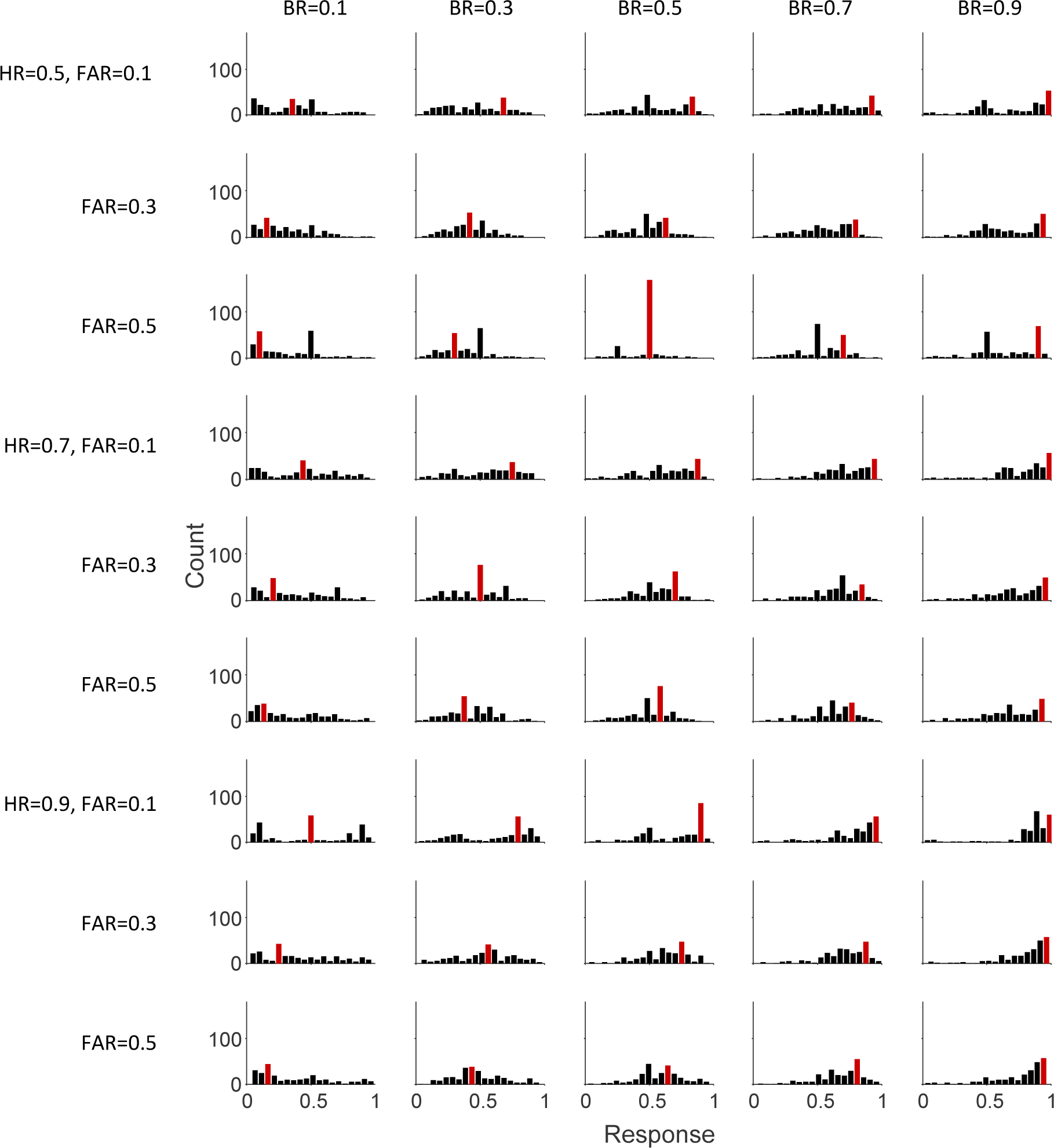
Distribution of participant responses in the main experiment. Histograms of participant responses on each of the 45 items in the main experiment, pooled across all four conditions. Each of the 45 items had of a unique combination of base rate (BR), hit rate (HR), and false-alarm rate (FAR) values. The red bar indicates the bin with the correct answer.

### Robustness check: re-analysis of base-rate sensitivity after excluding participants with extreme response times

Some of the participants in our experiment were suspiciously fast or slow. The median response time of the fastest participant was only 3.4 seconds, which seems barely enough to read the information on the screen, let alone to provide a serious response. On the other extreme, the longest participant had a median response time of 69 seconds, which suggests that external resources may have been used to solve the task. To verify that the individual differences in base-rate sensitivity were not solely due to participants with extreme responses times, we reanalysed the data with inclusion of only participants with median response times between 5 and 20 seconds, which corresponds to roughly 75% of the data. Both the distribution of sensitivity values (Figure A4, top) and the correlation between median response times and sensitivity index (*r* = 0.18, *p* = 0.02) were very similar as in the original analyses. The same holds when reducing the data further to include only the 50% most central median response times, albeit with a noisier pattern in the sensitivity distribution (Figure A4, bottom) and a further decrease in the strength of the statistical evidence for a correlation (*r* = 0.17, *p* = 0.08).

**Figure A4.**
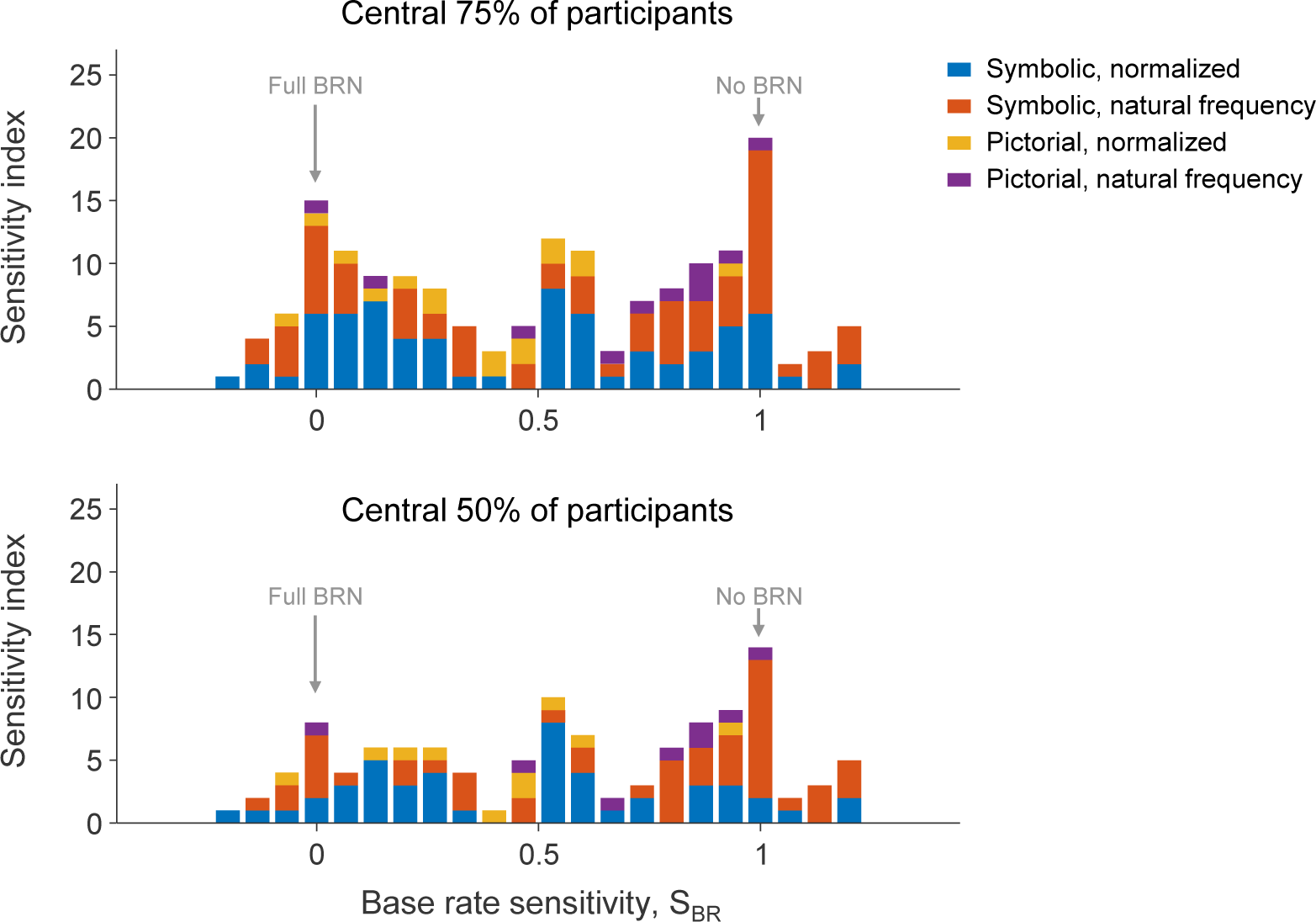
Base-rate sensitivity index distributions after filtering out participants with extreme response times. Top: the distribution of sensitivity indices for the central 75% of participants based on their median response times (12.5% percentile = 5.6 sec; 87.5% percentile = 24 sec). Bottom: the distribution for the central 50% of participants (25% percentile = 7.2 sec; 75% percentile = 17 sec).

### Comparison of predictions between the Linear-Additive model and the Bayesian mode

To examine if the Linear-Additive model and Bayesian model represent mathematically distinctive decision strategies, we scattered the predictions of both models against each other. While the results show a high level of correlation (Figure A5), they make quite different predictions on a large number of trials. This suggests that they implement mathematically distinct strategies.

**Figure A5.**
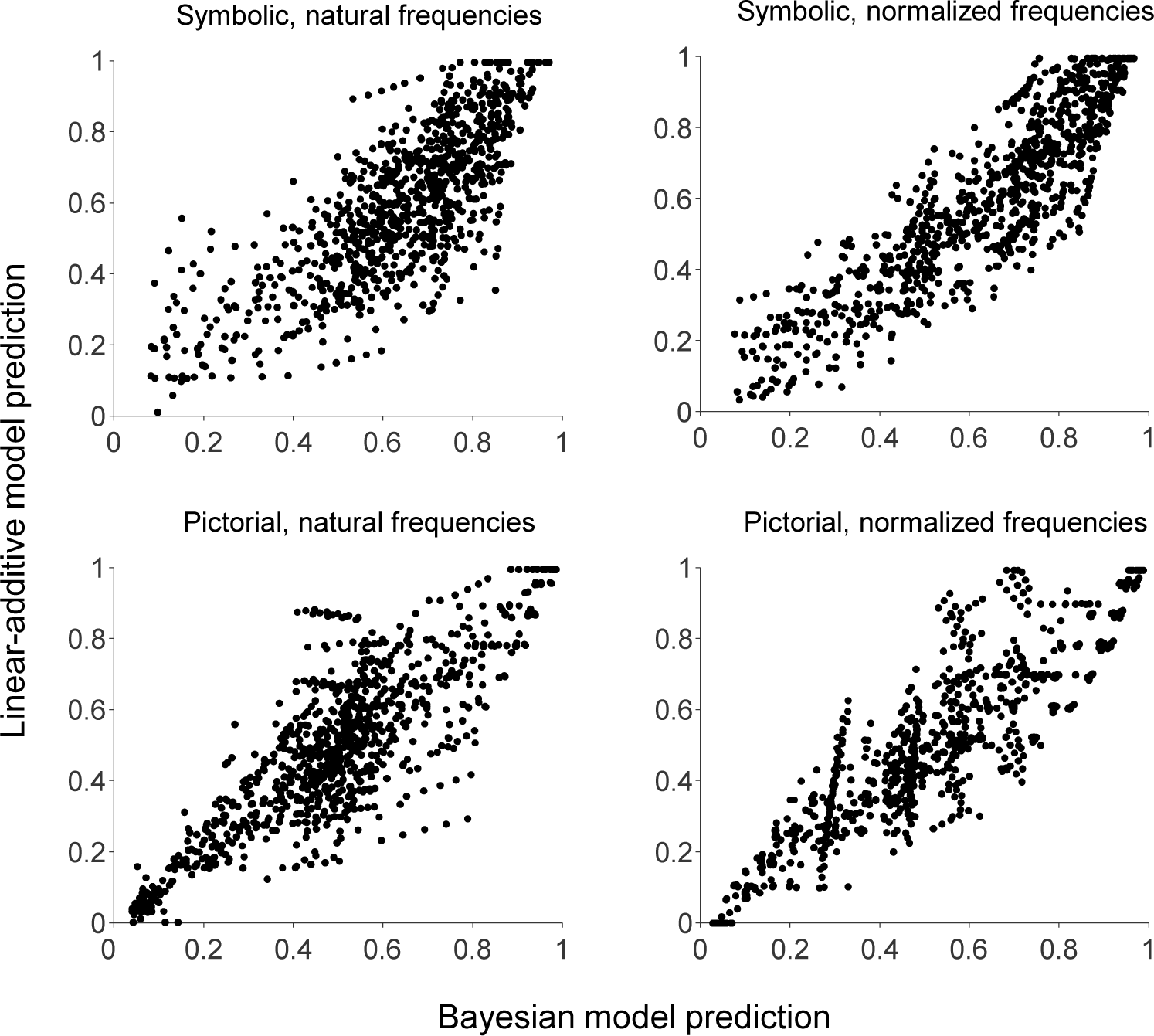
A comparison of the predictions by the Bayesian model and Linear-Additive model under maximum-likelihood parameter values. Each point represents the predicted response under both models on one of the 48 experimental trials. No early or late noise was included in these predictions. To avoid clutter, predictions are shown for only the first ten participants in each condition. Even though the model predictions are correlated – as expected since both were fitted to follow the empirical responses – on many of the trials they make quite distinct predictions.

### Robustness check: model comparison with a different decision noise distribution

To verify that the main modeling results do not critically depend on the specific choice for the decision noise distribution, we reran the model comparison with a Beta noise distribution, in which the mean on each trial was fixed to the model’s predicted response and the variance was fitted as a free parameter. The results of this analysis were highly similar to that of the main analysis (Figure A6).

**Figure A6.**
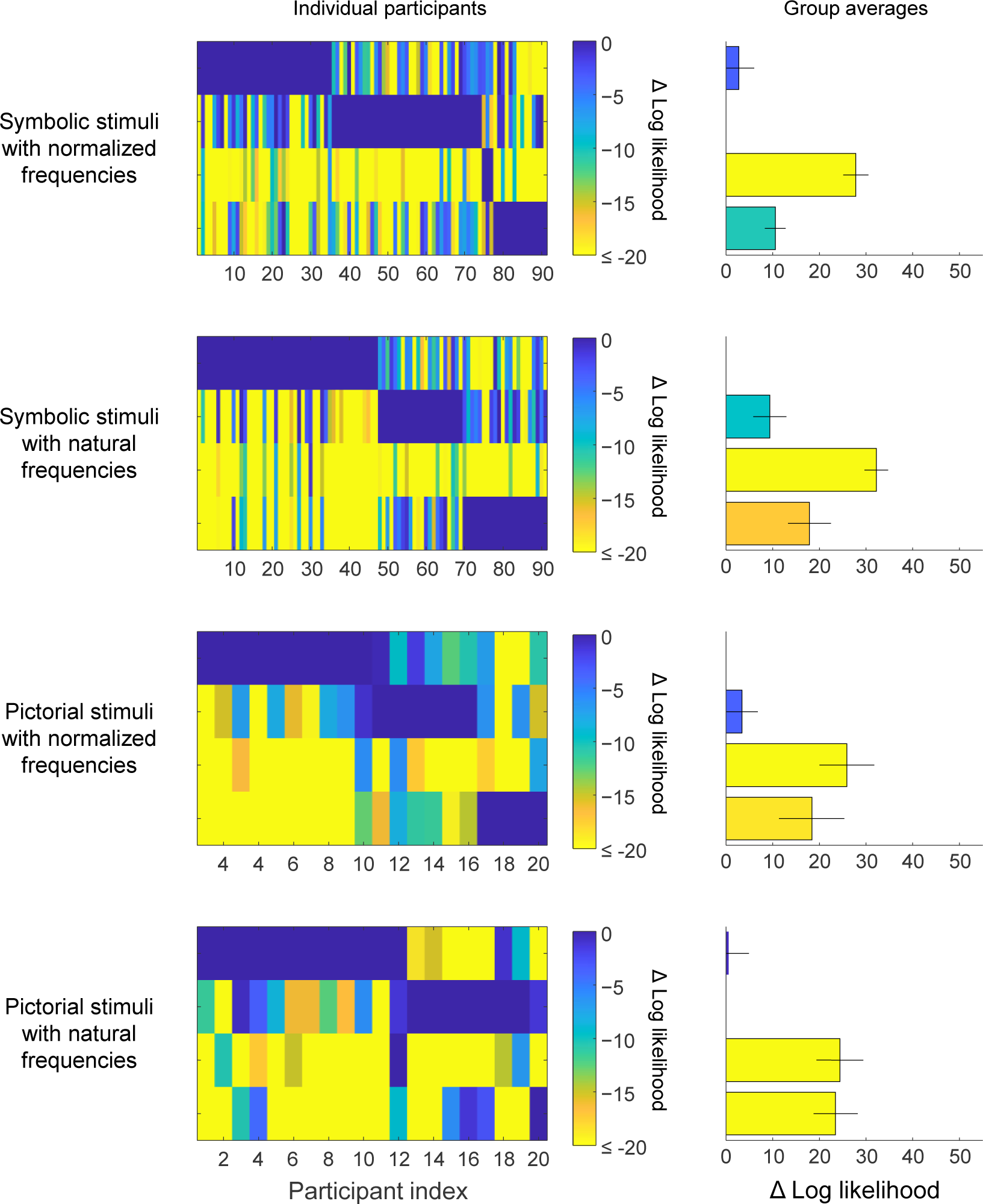
Model comparison results using an Beta distribution for decision noise. The model comparison results under this noise distribution look almost identical to those obtained using a Gaussian distribution on the log-odds model predictions (Figures 4-7 in the Main text).

### Simulation results 1: how well do the heuristic models approximate the optimal strategy?

To assess the ecological plausibility of the three non-Bayesian models, we examined how well they were able to approximate the Bayes-optimal responses. To this end, we optimized their parameters with respect to minimizing the root mean squared error (*RMSE*) between their predicted responses and the correct (Bayesian) responses on the 45 trials in our experimental task. Under these optimal parameters, all three models perform reliably better than a randomly guessing observer and an observer who always responds 0.50 (Figure A7A). This suggests that they are – in principle – viable strategies from an ecological perspective. The optimized linear additive model consistently outperforms the two heuristic models and has a relatively small error compared to a randomly responding observer and an observer who always responds 0.50. The optimal weights in the linear-additive model are *w*_BR_=1.11, *w*_HR_=0.52, and *w*_FAR_=−0.71, which means that the best performance is achieved by giving most weight to the base rate and least (in absolute terms) to the hit rate. According to the criterion estimates in the lexicographic model, the best possible performance is achieved when always considering the BR to be informative and to consider the HR and FAR informative when they deviate by more than approximately 0.30 from 0.50.

**Figure A7.**
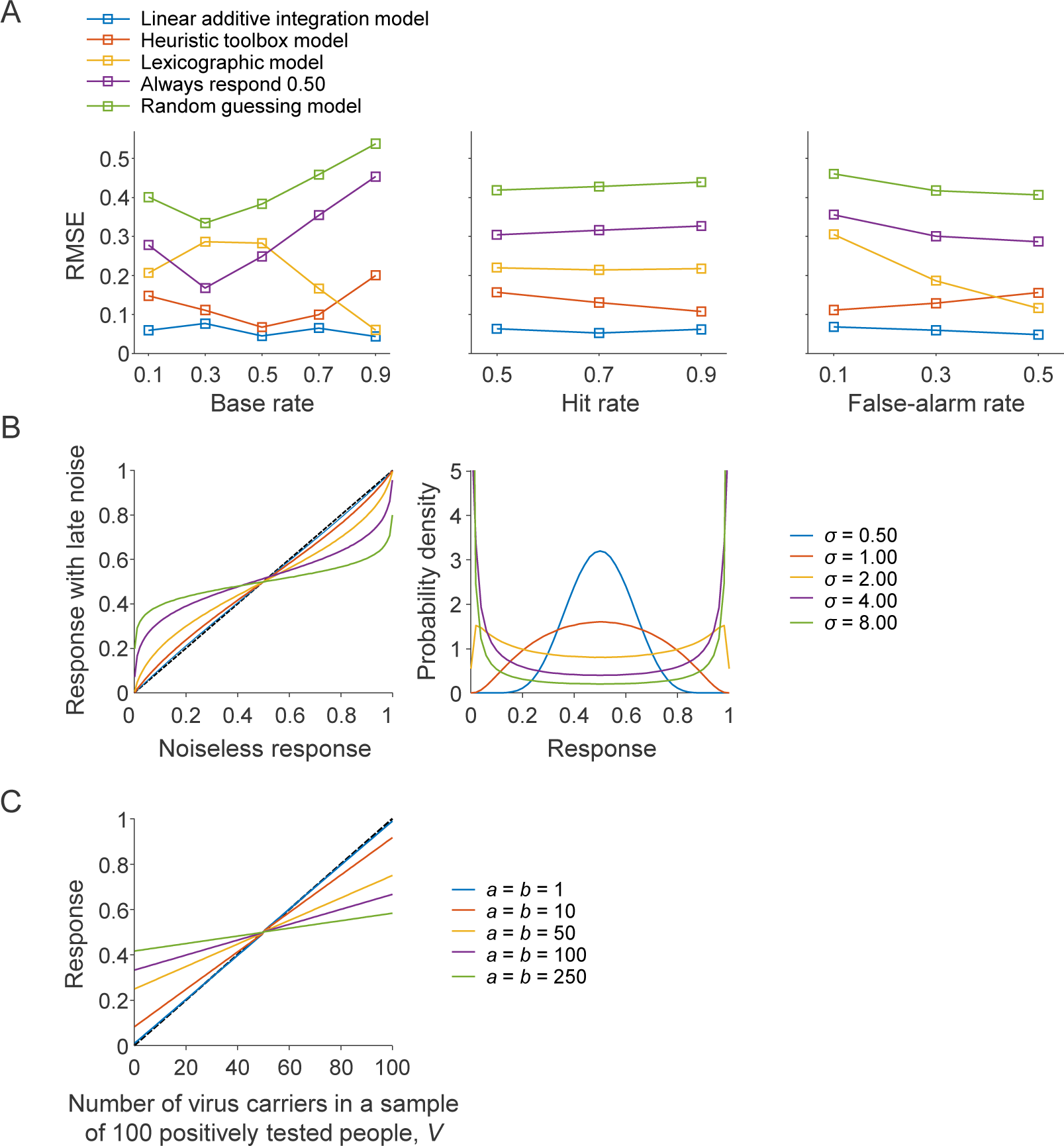
Analysis of model predictions. (A) Accuracy of the three non-Bayesian models on our experimental task, after optimizing their parameters with respect to minimization of the root mean squared error (RMSE). All models perform better than a randomly guessing observer (green) or an observer who responds 0.50 on each trial (purple), but with clear differences between them. (B) Regressive effects of late noise. Left: Mean response as a function of the decision before adding noise, for five different noise levels. Right: response distributions for the same five noise levels as in the left panel, for a trial in which the decision before adding noise was 0.50. Note that in order to obtain reasonably strong regression effects (yellow, purple, green in the left plot), noise levels are required that are so high that individual responses will have little connection with the initial decision. (C) Regressive effects of uninformative priors in the Bayesian model. The plot shows simulated responses for the medical diagnosis task in which the Bayesian observer is presented with the information that “*V* out of 100 positively tested people carried the virus”, under 5 different priors (different colors). In each case, the prior is “uninformative” (*a*=*b*). The stronger the prior (i.e., the higher the pseudo-counts), the stronger the response regresses towards 0.50.

It is notable how well the heuristic toolbox model performs, given that it has no free parameters. This shows that heuristic-based strategies can indeed be highly accurate, if participants know which heuristic to choose in any given situation. It is also striking how poorly the Lexicographic model performs, given that it has one more free parameter than the Linear-Additive model and four more than the Heuristic Toolbox model.

### Simulation results 2: regressive effects due to noise and priors

In probability judgment tasks it is often found that human responses are biased towards 0.50. Two different mechanisms have been proposed to explain this bias, both of which appear in the models that we test here. First, it has been found that certain forms of noise can drive the average response towards 0.50 (Costello & Watts, 2014, 2016, 2017, 2018, 2019). We find that the “late noise” included in the models tested here indeed causes this kind of bias (Figure A7B, left panel). Importantly, however, the effects are negligible unless noise levels are so high that the predicted responses are almost completely random (Figure A7B right panel). Another way to explain regressive effects in probability estimation tasks is to assume that participants have a prior belief in central values or, similarly, against extreme values (Zhu et al., 2020). Indeed, we find that the Bayesian model produces regressive effects when it has an “uninformative” prior (i.e., *a*=*b*), with the strength of the effect depending on the strength of the prior (Figure A7C).

An important difference, however, is that the Bayesian model does not need any form of noise to produce these effects. Hence, while noise and priors can both explain regressive effects in *average* responses, the two proposals make very distinct *trial-to-trial* predictions.

## Footnotes

1 For notational convenience, we will refer to this as “natural frequencies” throughout the rest of the paper.

2 The data in the present study show the same pattern (Figure 9)

3 As a robustness check, we also fitted the models using a Beta noise distribution, which gave similar results (see Results). Hence, our modeling results do not critically depend on the shape of the chosen noise distribution.

4 All reported Bayesian statistics were obtained using the default prior settings in JASP.

5 The different values of the base rate, hit rate and false alarm rate were entered as repeated measures factors and individual subject responses as dependent variable.

6 Note that a sensitivity of 1 does not necessary mean that the participant performed perfectly in accordance with Bayes’ theorem, because it is possible to produce incorrect responses while still adjusting the responses correctly to changes in the base rate (see Figure 3E for an example). Likewise, a BR sensitivity smaller than 1 does not necessarily mean that a participant was non-Bayesian: it could be a Bayesian observer with a prior belief about base rates. While the sensitivity index is useful for quantifying how strongly participants reacted to changes in the base rate, care should be taken when drawing conclusions about underlying cognitive mechanisms from this index.

7 A classic illustration of the divergence of the predictions by these two sorts of models is that the interaction plots for multiplicative models (like the Bayesian model) should yield fan patterns with diverging lines, whereas linear additive integration should produce parallel lines (see, e.g., the work by Norman H. Anderson (1996). In practice, when applied to probability, where all predictions have to be truncated to the 0 to 1 probability interval, these plots become messy and the complex patterns are better evaluated formally in terms of model fit than by testing specific interaction terms or visual inspection, and this is the line pursued here.

8 If all participants are included, *Mdn* = 0.67, *IQR* = 0.53 for the Bayesian model and *Mdn* = 0.55, *IQR* = 0.35 for the linear model.

